# The liverwort *Marchantia polymorpha* operates a depolarization-activated Slowpoke (SLO) K^+^ channel that recognises pH changes in the environment

**DOI:** 10.1101/2021.06.01.446568

**Authors:** Frances C. Sussmilch, Jennifer Böhm, Guido Gessner, Tobias Maierhofer, Thomas D. Müller, Stefan H. Heinemann, Dirk Becker, Rainer Hedrich

## Abstract

Voltage-dependent ion channels are a prerequisite for cellular excitability and electrical communication – important traits for multicellular organisms to thrive in a changeable terrestrial environment. Based on their presence in extant embryophytes and closely-related green algae, the first plants to survive on land likely possessed genes encoding channels with homology to large-conductance calcium-activated K^+^ channels (BK channels from the Slo family) in addition to primary voltage-gated potassium channels from the plant VG-type family (Shaker or K_v_ channels). While the function and gating of Shaker channels has been characterised in flowering plants, so far knowledge of BK channels has been limited to animal models. In humans, BK-mediated K^+^ efflux has a critical role in sperm motility and membrane polarisation to enable fertilisation. In the liverwort *Marchantia polymorpha*, the *MpBK2a* channel gene is most highly expressed in male reproductive tissue, suggesting that these channels may function in sexual reproduction. We characterised MpBK2a channels and found them to be strongly K^+^-selective, outward-rectifying, 80-pS channels capable of repolarising the membrane after stimulus-dependent depolarisation. In contrast to its animal counterpart, MpBK2a is insensitive to cytoplasmic Ca^2+^ variations but effectively gated by pH changes. Given that this plant BK channel is active even in the presence of trace amounts of external K^+^ and at low pH, the liverwort channel could have stabilised the membrane potential under stressful pre-historic conditions including nutrient-depleted and acid environments as early plant pioneers conquered land.

## Introduction

Around 500 Mya, the first plants conquered dry land, gradually transforming it from a barren environment – where previously only bacteria and fungi had survived – to an environment that could sustain diverse life forms, including animals. The ancestors of land plants were likely streptophyte algae that developed important adaptations necessary to survive terrestrial conditions, including sturdy cell walls, protection against abiotic stresses, and elaborate cell-to-cell signalling pathways to enable co-ordinated, whole-body responses to environmental changes [1–3]. Extant bryophytes, such as the simple thalloid liverwort *Marchantia polymorpha*, offer insight into the earliest group of plants that thrived on land [4].

Similar to animals, Ca^2+^ flux and changes in the plasma membrane electrical potential, both mediated by ion channels, are central to electrical signalling for cell-to-cell communication in plants. The “green” action potential (AP) is shaped by Cl^−^ efflux-based depolarisation and repolarisation via K^+^ efflux, which resets the cell for the next electrical event [5]. SLAC1-type and QUAC1-type anion efflux channels are found throughout land plants [6–9]. Voltage-gated, outward-rectifying K^+^ channels, which share some structural and sequence features with the Shaker-type (K_v_ family) channels that shape animal APs, have also been characterised in flowering plants, including GORK and SKOR in *Arabidopsis thaliana* [10, 11]. Such channels are present in all major land plant groups except mosses [9, 12–14]. An additional K^+^ channel type with similarity to the animal Slo family, which includes BK (large-conductance Ca^2+^- and voltage-activated channels, also named Slo1 or KCa1.1, encoded by *KCNMA*1 in humans), Slo2 (Slick/Slack) and Slo3, has also been found in the genomes of non-flowering plant clades [9, 12]. Animal *Slo* genes each encode α subunits for homotetrameric K^+^ channels that have high single-channel conductance (up to 300 pS for BK) and are activated by concerted membrane depolarisation and elevated intracellular Ca^2+^ concentrations. These Slo channels are important for various aspects of normal animal physiology including electrical signalling in neurons and smooth muscle, and male fertility through the regulation of sperm motility and competence for fertilisation [15–17]. Currently available genome data of flowering plants suggest they encode only Shaker-type voltage-dependent K^+^ channels. In contrast, the *Marchantia* genome also encodes voltage-dependent K^+^ channels of the BK type. Here we report the properties of a *Marchantia polymorpha* BK channel that may provide for K^+^ homeostasis and electrical signalling in non-flowering land plants.

## RESULTS

### The evolution of plant BK channels

To identify members of the BK/Slo channel family in plant lineages, we used an HMM profile built from characterised animal Slo channel α subunits. We found a strongly-supported clade of BK channels encoded within the genomes of diverse non-flowering plant groups, ranging from green algae to gymnosperms (**Figures 1A and S1**; **Table S1**). Homologous genes, though, were not present in angiosperms examined. In addition to plants and animals, we also identified the presence of a Slo1/3-like channel gene in an amoeba and Slo2-like channel genes in choanoflagellate, ichthyosporea and fungal species, suggesting that the two main groups of animal Slo channels arose from an ancient duplication, prior to divergence of these eukaryote clades.

**Figure 1.**
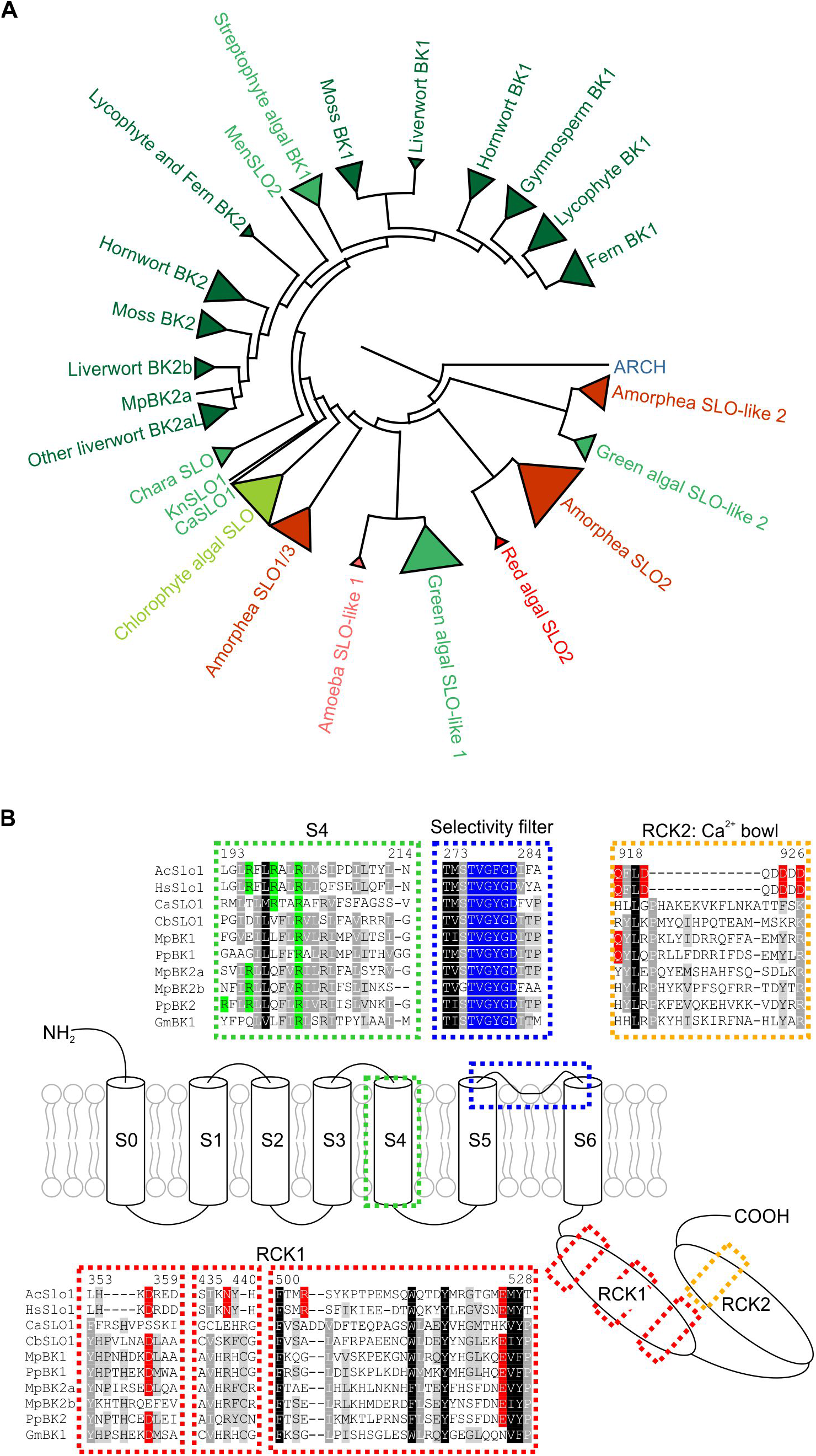
Plants encode BK channels, similar to other eukaryotes. (A) A condensed phylogeny of the eukaryote Slo family, rooted to a related archaeon gene. The fully expanded tree (Figure S1) is displayed with collapsed subclades (triangles). See Table S1 for full species and gene details. (B) Schematic representation of the domain architecture of animal BK channels, based on AcSlo1 [27], and selected regions of aligned animal (Ac, *Aplysia californica*; Hs, *Homo sapiens*) and plant including algal (Ca, *Chlorokybus atmophyticus*; Cb, *Chara braunii*), bryophyte (Mp, *Marchantia polymorpha*; Pp, *Physcomitrella patens*) and gymnosperm (Gm, *Gnetum montanum*) BK sequences. Displayed regions include the S4 transmembrane domain showing conserved charged amino acids linked to voltage dependency (green), the conserved K^+^ selectivity filter in the pore (dark blue), and residues critical for Ca^2+^ sensitivity in the RCK1 domain and Ca^2+^ bowl in the RCK2 domain (red) of animal BK proteins. Numbers indicate sequence position in AcSlo1.

The majority of bryophyte, lycophyte and fern species queried possessed two or more BK genes, but only a single BK gene was identified in each gymnosperm species examined (**Figure S1**), which may reflect a reduction in the selection pressure to preserve multiple BK genes in seed plants. One feature that many of these BK-harbouring plant species share is flagellated cells, which are not produced by angiosperms. Some streptophyte algae, including *Chlorokybus atmophyticus* and *Klebsormidium nitens* have flagellated zoospores [18, 19], while the Charophyceae (including *Chara*), bryophytes, ferns and some gymnosperms (including *Ginkgo*), have multi-flagellated spermatozoa, which typically move through the external environment to reach non-motile female gametes.

In gymnosperms that have retained flagellated sperm, such as *Ginkgo*, these are required to swim only a short distance in a controlled environment, as male genetic material is delivered inside female reproductive tissues in the robust form of pollen. In angiosperms, which lack BK genes, immotile sperm cells are delivered directly to the ovule by pollen tube growth, and – in this controlled environment, where external potassium is available – pollen-expressed outward-rectifying Shaker channels, including SKOR in Arabidopsis [20, 21], may account for voltage-dependent K^+^ efflux from the pollen tube [22].

Like the structurally related, voltage-gated Shaker channels, BK channels assemble into functional tetramers, with each pore-forming α-subunit comprising the following functional domains: the pore-gate domain (PGD, formed by transmembrane helices S5-S6 and harbouring the selectivity filter and the ion conducting pathway) and the voltage-sensing domain (VSD, formed by transmembrane helices S1-S4; **Figure 1B and Figure S2**). This VSD contains a motif of arginine residues interspaced with hydrophobic residues – RxxR – that is conserved between voltage-sensing cation channels [23]. In contrast to Shaker channels, BK channels typically exhibit an extra S0 helix that positions the BK N-terminus extracellularly. Finally, a ligand-sensing, cytosolic tail domain (CTD) comprising the so-called regulator of conductance for K^+^ (RCK) domains locates downstream of the BK transmembrane moiety [24–26]. In animal BK channels, the intracellular RCK gating ring typically harbours the Ca^2+^ binding sites, communicating cytosolic Ca^2+^ changes to voltage-dependent gating of the channels.

To study the possible role of plant BK channels, we selected *Marchantia polymorpha* as a land plant model for further characterisation of channel properties because this simple thalloid liverwort offers the availability of genome and RNA-seq data [4]. Structural modelling of Marchantia BK2a using available cryo-electron microscopy-derived structures of Slo1 channels from Aplysia and Human [24, 27] allowed us to get first insight into the 3D architecture of the plant BK channel. Despite overall low sequence conservation (~23% identity), the MpBK2a amino acid sequence could be mapped satisfactorily onto the AcSlo1 and hSlo1 structures (**Figure S3**). In line with the notion that no signal peptide was predicted for the MpBK2a protein, a S0 transmembrane helix was not properly assigned in the overall protein alignment with animal BK channel proteins. S0, however, was part of the structural model, implying that the N-terminus of the liverwort BK channel might locate to the extracellular site as reported for its animal counterparts [28, 29]. The ion channel moiety comprising the voltage-sensing domain (VSD, S1-S4) and the pore-gate domain (PGD, S5-S6) properly aligned with both Slo1 structures demonstrating that the transmembrane, ion-conducting part of the channel is highly conserved between animals and plants (**Figure 1B**). Downstream of the transmembrane domain (TMD), the existence of two RCK domains was confirmed by structural prediction in the CTD of the plant BK channel protein. In comparison to its animal counterparts, the amino acid stretch constituting the RCK1-RCK2 linker is significantly extended, implying an altered interaction and/or spatial architecture of these regulatory domains in the Marchantia BK channel. Further, key residues responsible for the coordination of Ca^2+^ in the animal BK channel RCK domains [27, 30] are poorly conserved in MpBK2a, challenging the idea of a possible allosteric regulation of MpBK channel gating by intracellular divalent cations and voltage.

### The *MpBK2a* gene encodes a plasma membrane K^+^ channel activated by membrane depolarisation

Available RNA-seq data indicates that Marchantia *BK* channel genes are most highly-expressed in male reproductive tissue [4, 31], suggesting that these channels may function in sexual reproduction, similar to mammalian BK and Slo3 channels. Among the three BK channel genes in the Marchantia genome, *MpBK2a* exhibited the broadest expression pattern (https://evorepro.sbs.ntu.edu.sg) [32] and – besides male reproductive tissue – was also expressed in vegetative thallus tissue (**Figure 2A**).

**Figure 2.**
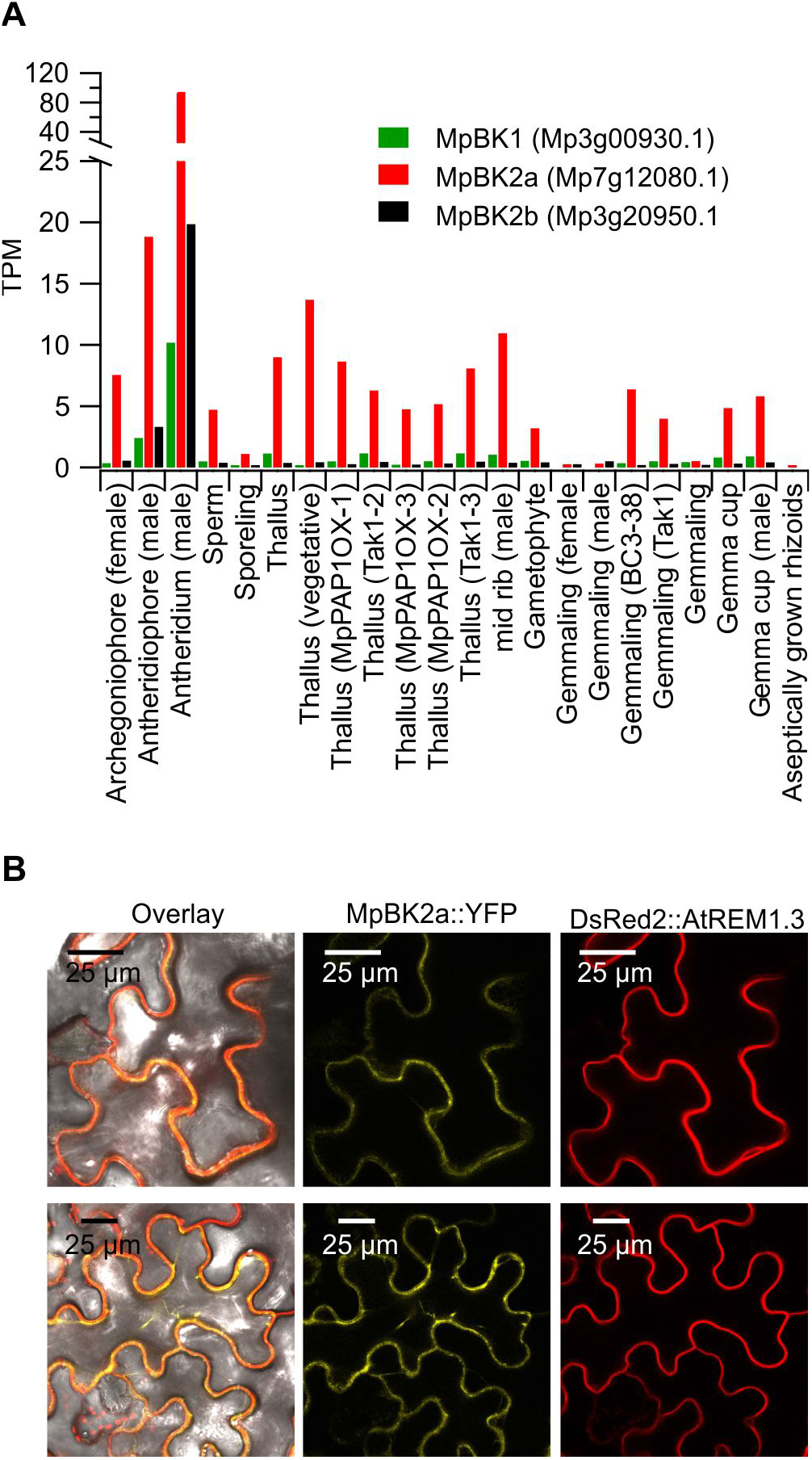
Transcriptional profile of *Marchantia polymorpha* MpBK2a and its localisation to the plasma membrane. (A) Transcriptional profile of *Marchantia polymorpha* BK channel expression. Data were obtained according to available RNA-Seq analyses from CoNekT (https://evorepro.sbs.ntu.edu.sg/). (B) MpBK2a fused C-terminally to YFP transient transformed into *N. benthamiana* cells and imaged three days post infiltration (middle panel). DsRed2::AtREM1.3 was used as plasma membrane marker (right panel). The overlay of cells co-expressing MpBK2a::YFP and DsRed2::AtREM1.3 (left panel) indicates localisation of MpBK2a at the plasma membrane.

To elucidate the subcellular localisation of the Marchantia BK2a channel, we employed leaf cells of *Nicotiana benthamiana* cells as transient expression systems [33]. Upon co-infiltration with the plasma membrane marker DsRed2::AtREM1.3 [34], we observed co-localisation and thus plasma membrane targeting for MpBK2a using confocal laser scanning microscopy (**Figure 2B**), suggesting a functional role in cytosolic K^+^ homeostasis and/or electrical signaling.

This prompted us to express the Marchantia BK channels heterologously in *Xenopus laevis* oocytes asking the question of how do plant BKs respond to voltage changes. We selected the most dominant BK out of three present in *Marchantia polymorpha* (MpBK2a; Mapoly0003s0221; Mp7g12080)[4] for electrophysiological characterisation. To study the voltage-dependent features of MpBK2a, we injected cRNA into *Xenopus* oocytes and measured ion currents using the Two Electrode Voltage Clamp (TEVC) technique. Under essentially symmetric potassium conditions, i.e. 100 mM K^+^ in the bath solution, voltage pulses from −40 to 100 mV elicited outward currents with a slow activation kinetic that reached steady-state within 1 s upon membrane depolarisation in oocytes expressing MpBK2a (**Figure 3A and B**). With 100 mM K^+^ in the bath, tail currents reversed close to 0 mV. Upon replacement of K^+^ in the bath solution by equimolar concentrations of either Na^+^ or Li^+^, steady-state outward currents remained largely unchanged. Thus, in the absence of external K^+^ channel gating or activation kinetics appeared unaffected (**Figure 3A and 3C**). Under these conditions, however, inward tail currents essentially disappeared, indicating that K^+^ but not Na^+^ or Li^+^ are conducted through open BK channels. In addition, voltage-dependent gating and activation kinetics of MpBK2a was not affected by extracellular K^+^ (**Figure 3C**). Plant cells typically face extracellular, cell-wall K^+^ concentrations in the low millimolar range. Increasing the driving force for K^+^ by lowering [K^+^]_ext_ from 100 mM and 10 mM to finally 1 mM resulted in outward steady-state currents of slightly increased amplitude, shifted the reversal potential of the MpBK2a tail currents in a K^+^ dependent manner (**Figure S4A**) but did not affect the voltage-dependence of channel activation (**Figure 3D and E**). Taken together, this shows that MpBK2a constitutes a depolarisation-gated, outward-rectifying K^+^ channel but — in contrast to plant Shaker-type K^+^ efflux channels — appears to lack a regulatory site for extracellular K^+^ [35].

**Figure 3.**
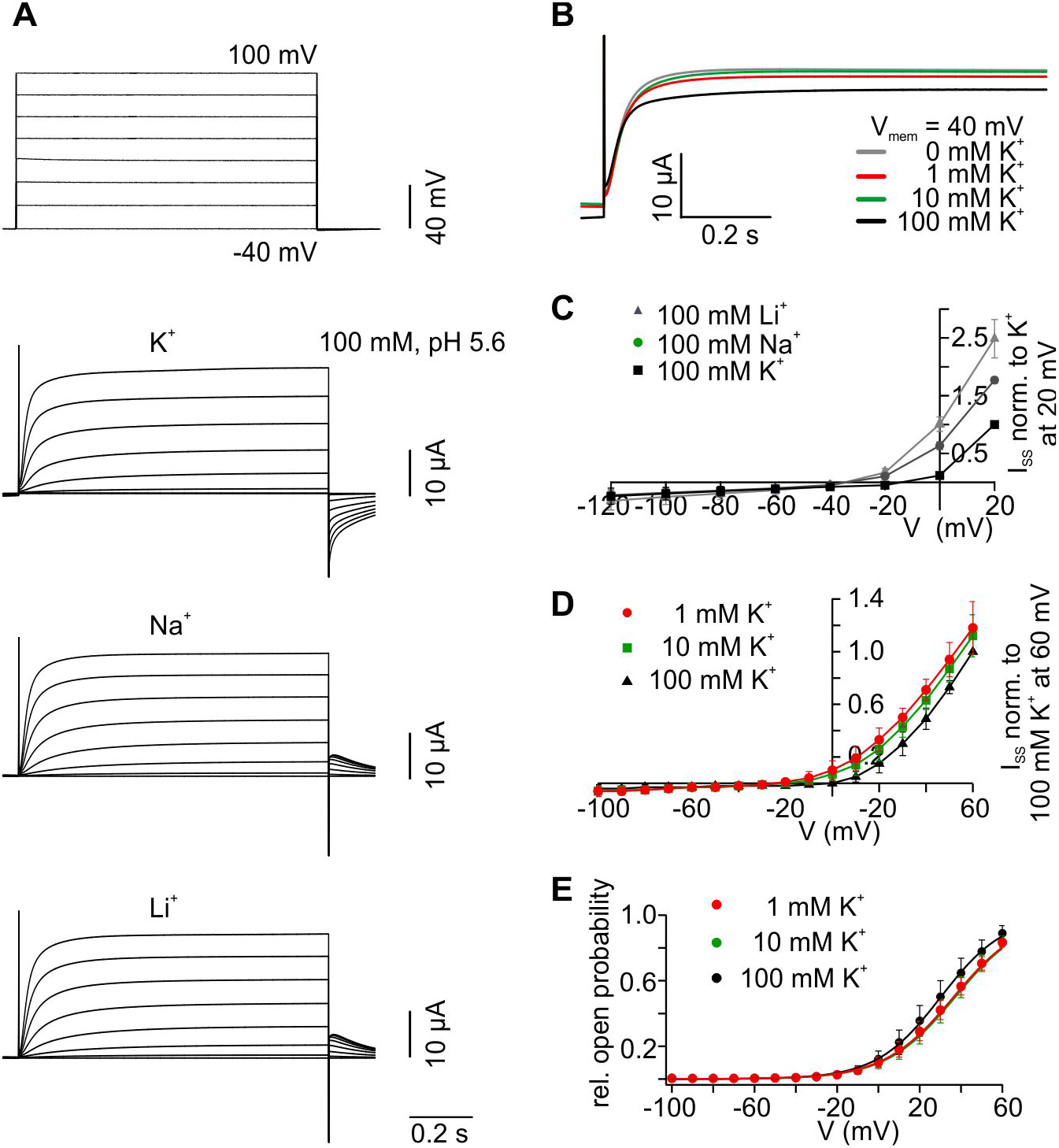
MpBK2a is a depolarisation-activated K^+^ channel. (A) Original traces of outward-rectifying currents measured by TEVC in a representative MpBK2a-expressing oocyte perfused with 100 mM KCl, NaCl and LiCl solutions (pH 5.6) upon depolarising voltage steps between −40 mV (V_H_) to 100 mV in 20 mV increments (pulse protocol at the top). Negative I_tail_ are consistent with cation influx and are seen only when oocytes are perfused with solutions containing K^+^, indicating that MpBK2a is highly K^+^ selective. All current traces are shown at the same scale as indicated. (B) Representative current traces of MpBK2a-expressing oocyte at 40 mV in varying K^+^ concentrations. The slow activation kinetic is not affected by a decrease in the [K^+^]_ext_. (C) Steady-state currents (I_SS_) recorded in 100 mM K^+^, Na^+^ and Li^+^ (chloride-based solutions; pH 5.6) were plotted against the applied voltages. Note that I_SS_ are highest when potassium is not added to external solutions (n = 6, mean ± SD). (D) Steady-state currents mediated by MpBK2a in varying K^+^ concentrations remain largely unaffected by the external concentration change (n = 6, mean ± SD). (E) The relative open probability of MpBK2a is not dependent on external potassium concentration (V_0.5_ = 21.7 ± 8.7 mV at 100 mM K^+^; V_1/2_ = 35.0 ± 4.5 mV at 10 mM K^+^; V_1/2_ = 34.8 ± 3.8 mV at 1 mM K^+^). Relative open probability was determined from I_tail_ following a shift from test pulses in the voltage range −100 to 60 mV to a 50 mV pulse. Data points were fitted with a Boltzmann function (solid lines, n ≥ 4, mean ± SEM). Gluconate-based solutions were used to obtain the K^+^ concentrations shown.

For control over the ion content at the cytosolic face of the oocyte membrane (100 mM NaCl or 100 mM KCl; **Figure S4B and C**) we used macro patches of MpBK2a-expressing oocytes in the inside-out configuration with 100 mM K^+^ in the patch pipette. When perfusing the membrane patch with Na^+^-based buffer (100 mM), no outward currents could be measured in the whole voltage range explored (−60 to 80 mV; **Figure S4B and D**). However, triggered by a short pre-activation pulse (80 mV), macroscopic inward tail currents were monitored over a voltage range from −60 to 80 mV, associated with K^+^ fluxes from the pipette into the bath (**Figure S4E**). In contrast, when K^+^ was present in the bath (100 mM), macroscopic outward currents were recorded with kinetics and voltage dependencies reminiscent of whole-oocyte MpBK2a currents recorded in TEVC measurements (**Figure S4C and D**). Under these conditions, tail currents reversed around 0 mV (**Figure S4E).** In line with the presence of the conserved K^+^ selectivity filter (TVGYGD) in the pore loop of MpBK2a (**Figure 1B**) and the TEVC recordings in *Xenopus* oocytes, this observation supports the notion that MpBK2a is K^+^ selective (**Figure S4A**).

We also performed patch-clamp analyses with MpBK2a-expressing HEK293T cells to further elucidate the biophysical properties of the Marchantia BK channel. In whole-cell recordings we essentially confirmed observations from oocyte measurements, indicating that the voltage-dependent K^+^ channel features of the MpBK2a channel do not depend on the heterologous expression system (**Figure 4**). In inside-out patches from HEK293T cells with symmetrical 100 mM K^+^ solutions, the MpBK2a single-channel conductance in the inward and outward direction of about 80 and 40 pS was determined, respectively (**Figure 4**). Thus, the single-channel conductance of MpBK2a is about 2-4 fold higher compared to plant Shaker-type K^+^_out_ channels [36, 37] but significantly smaller compared to animal Slo1 channels with up to 300 pS [38]. Neither the typical animal BK activators NS1619, NS19504, and NS11021 nor the BK inhibitors iberiotoxin, paxilline and penitrem A [25] markedly affected MpBK2a-mediated currents indicating that the specific pharmacological features of animal BK channels are not conserved in MpBK2a (**Figure S5**). Typical for a voltage-dependent, K^+^-selective plant channel inhibition of current amplitude was observed on oocytes in TEVC experiments in response to external perfusion with millimolar concentrations of the K^+^ channel blockers tetraethylammonium (TEA^+^) and Ba^2+^ (to a lesser extent) with maximal currents reduced by 40% (10 mM TEA^+^) and 20% (10 mM Ba^2+^), respectively (**Figure S4F**).

**Figure 4.**
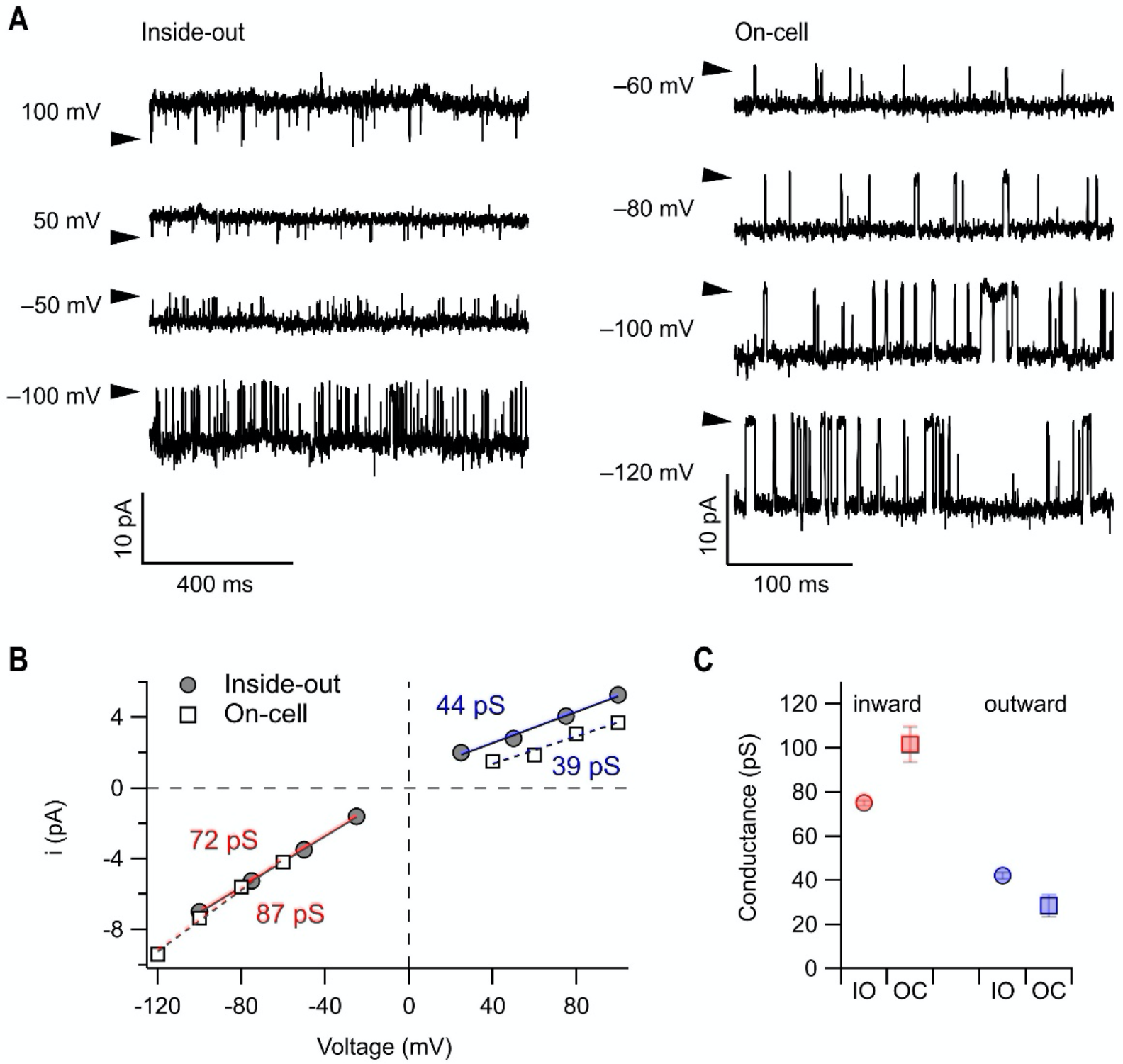
Single-channel recordings in HEK293T cells. (A) Representative single-channel current traces for the indicated voltages in inside-out mode (left) and in the on-cell configuration (right). The arrowhead mark closed channel current levels. (B) Mean open-channel current amplitudes as a function of voltages from the experiments shown in (A). The superimposed lines indicate linear fits to estimate the single-channel conductance in the inward (red) and outward (blue) direction. (C) Mean single-channel conductance. Data points are means ± SEM (n = 4 experiments each); IO, inside-out; OC, whole-cell. Solutions, concentrations in mM: IO, symmetrical 140 KCl, 10 EGTA, pH 6.0 (KOH); OC, pipette, 100 KCl, 50 NaCl, 2 MgCl_2_, 2 CaCl_2_, 10 HEPES, pH 6.0 (NaOH).

### MpBK2a is sensitive to physiological pH changes but not to Ca^2+^

In animal BK channels, the RCK1 and RCK2 domains harbour residues critical for Ca^2+^ and Mg^2^ binding and activation of the K^+^ channel [26, 27, 39]. Whole-cell patch-clamp measurements of HEK293T cells expressing MpBK2a showed that increasing the intracellular Ca^2+^ concentration from nominally 0 to 100 μM did not increase the current amplitude or result in significant alteration in the half-activation potential (V_0.5_) (**Figure 5 and Figure S6**). Moreover, and in line with the low sequence conservation between plant and animal BK channels, neither cytosolic Mg^2+^ (2 mM) nor Na^+^ (10 mM) modulated MpBK2a currents, further distinguishing the plant BK channel from its animal counterparts of the Slo1 and Slo2 subfamily, respectively (**Figure 5**). MpBK2a-mediated steady-state currents decreased in the presence of cytosolic cations at membrane potentials positive of 60 mV (**Figure 5 and Figure S6**), indicative for a block by cytosolic cations. This demonstrates that the maximal MpBK2a activity does not require the elevation of cytoplasmic Ca^2+^ levels. Thus, in contrast to animal BKs, the liverwort BK channel is neither sensitive to the pharmacology nor to intracellular Ca^2+^ concentration changes that address animal BK regulatory sites.

**Figure 5:**
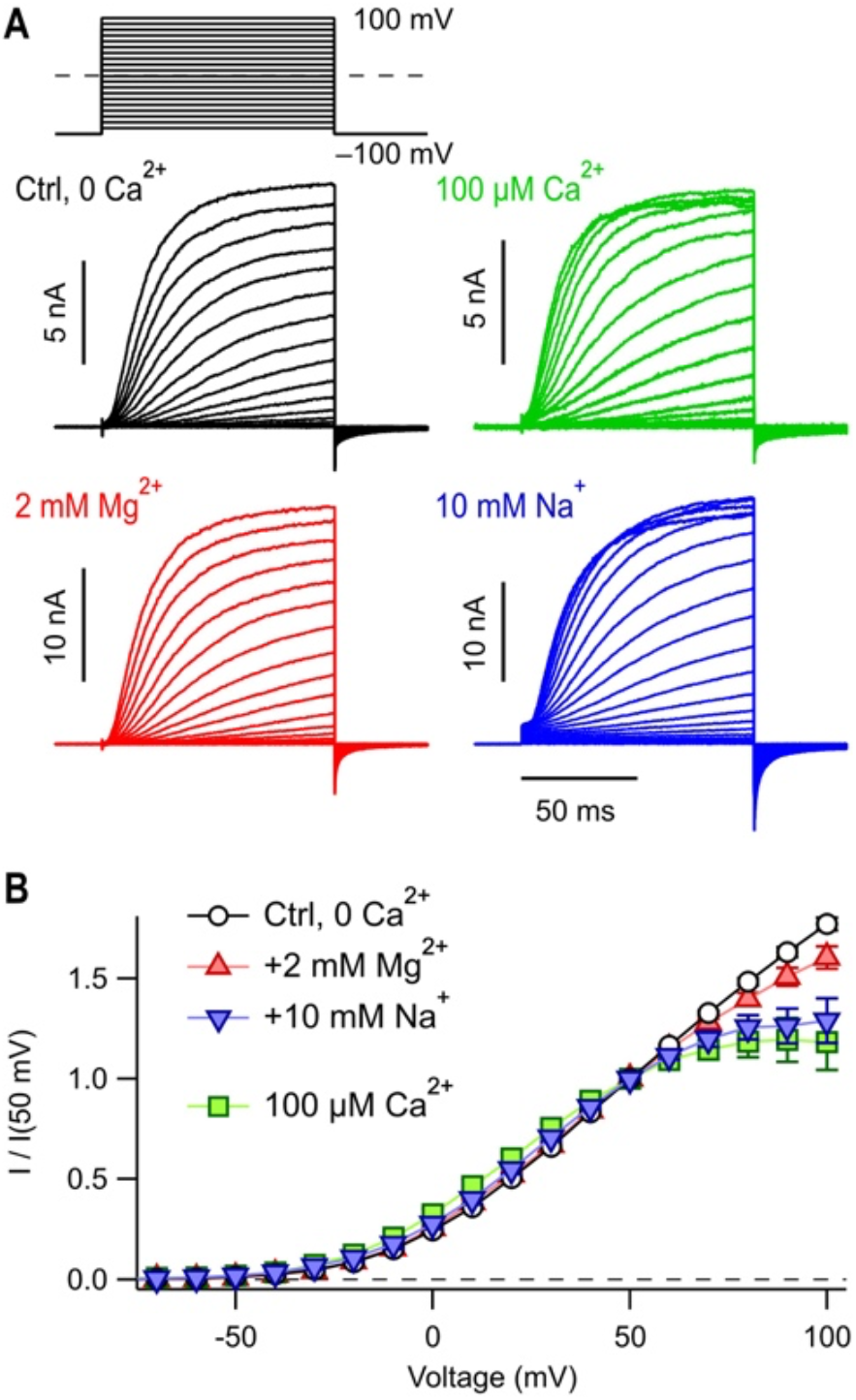
MpBK2a is not gated by intracellular Ca^2+^, Mg^2+^ or Na^+^. (A) Superimposed whole-cell current traces of MpBK2a expressed in HEK293T cells subjected to the pulse protocol indicated. Extracellular (bath) solution (in mM): 135 NaCl, 5 KCl, 2 MgCl_2_, 2 CaCl_2_, 10 HEPES, pH 7.4 (NaOH). Intracellular (pipette) solutions (in mM): Ctrl, 140 KCl, 10 EGTA, 10 HEPES, pH 7.4 (KOH); “Mg^2+^”, plus 2.56 MgCl_2_ yielding 2 mM free Mg^2+^; “Na^+^”, plus 10 NaCl; “100 μM Ca^2+^”, plus 100 μM CaCl_2_ without EGTA. (B) Maximal outward currents of data as in (A), normalised to the current at 50 mV. Data points are means ± SEM (n = 5-6). Straight lines connect data points for clarity.

As sessile organisms, plants are often faced with unstable environmental conditions such as nutrient availability and soil acidity. This was likely even more pronounced at the time plants conquered land. Furthermore, high affinity uptake of the macronutrient K^+^ is energised by an outward-directed H^+^ pump. That is why even in the plant body, the extracellular compartment, the apoplast, is generally acidic [40]. To answer the question of MpBK2a sensitivity towards the external proton concentration, we challenged MpBK2a with physiological pH variations [41, 42]. Using both TEVC in the oocyte expression system as well as patch clamp with HEK293T cells, we found that the voltage-dependent activation of the channel changed as a function of the external pH (**Figure 6 and S7**). In either expression system, MpBK2a-mediated outward steady-state currents increased upon extracellular alkalinisation (**Figure 6A and S7A**). Independent of the external K^+^ concentration, an increase in external pH by 1.5 units resulted in a prominent negative shift of the half-activation potential by about 50 mV **(Figure 6B and S7B**). In addition, external protons had a notable influence on the activation and deactivation kinetics of Marchantia BK2a, with activation slowed down and deactivation accelerated at acidic pH (**Figure 6D and E, Figure S7C**). In oocytes, the observed pH-dependent shift in activation potential, combined with slower channel closure, resulted in inward K^+^ currents at negative membrane potentials within the range of 0 to −60 mV when 100 mM K^+^ was applied externally (**Figure S7A**). Cytoplasmic acidification – a situation frequently observed in plant cells upon biotic and abiotic stress conditions – also affected voltage-dependent gating of MpBK2a, but compared to extracellular H^+^, shifted V_0.5_ to negative membrane potentials. Thus, intracellular H^+^ stimulates MpBK2a activity, favouring channel opening and resulting in increased steady-state currents at depolarising membrane potentials (**Figure 7**).

**Figure 6.**
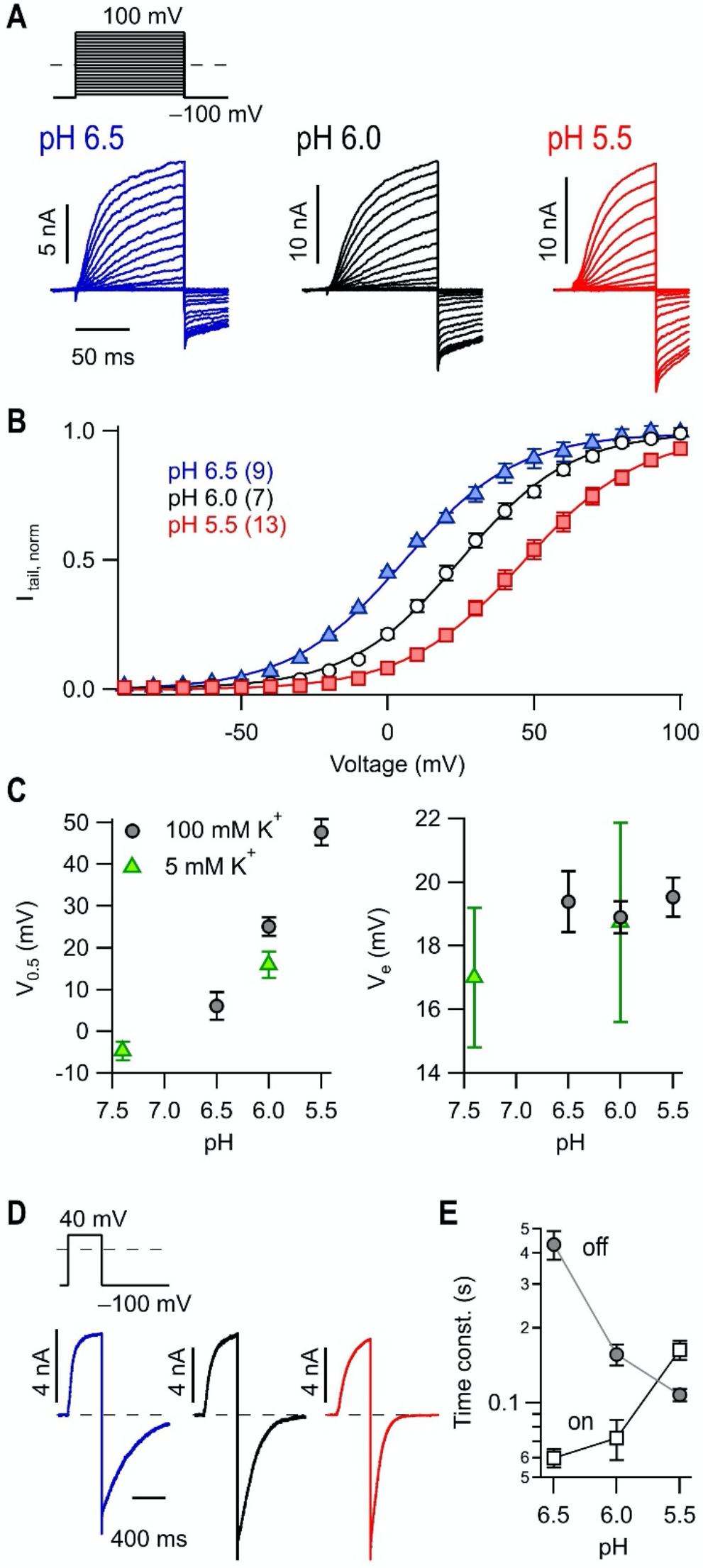
(A) MpBK2a gating depends on extracellular pH. (A) Pulse protocol and superposition of whole-cell current traces from HEK293T cells expressing MpBK2a with the indicated extracellular pH values. Solutions with symmetrical K^+^ concentrations were used to assess tail currents. (B) Mean maximal tail currents as a function of the prepulse potential with superimposed fits according to Boltzmann functions. Currents were normalised to the saturating currents derived from the fits. Data are means ± SEM (n in parentheses). (C) Results from the fits shown in (B) as a function of pH: *V*_0.5_, the half-maximal activation voltage; *V*_e_, the corresponding slope factor. (D) Current traces according to the indicated pulse protocol with extended time base to assess kinetics of activation and deactivation. pH values color-coded as in (A, B). (E) Mean time constants of activation (square) and deactivation (circles), determined from data as in (D) with single-exponential fits, as a function of pH. Data points are means ± SEM (n = in parentheses), connected by straight lines for clarity.

**Figure 7.**
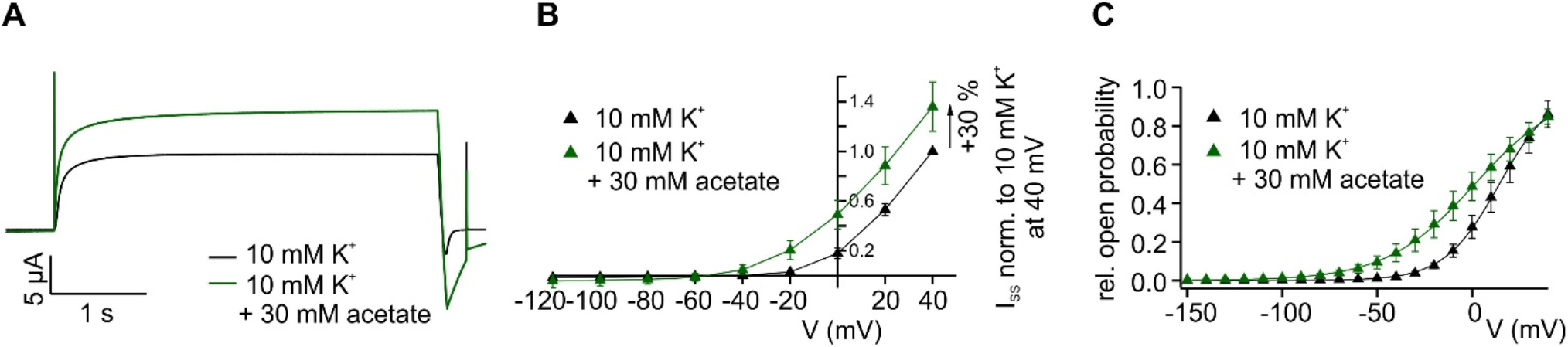
Cytoplasmic protons activate MpBK2a. (A) Representative current traces monitored by a voltage ramp of a MpBK2a-expressing oocytes in 10 mM K^+^ (black; pH 5.6) and after the internal acidification with 30 mM acetate (green; pH 5.6). (B) The I_SS_-voltage relation at voltages ranging from −120 to 40 mV revealed an increase in the outward current amplitude by 30% after perfusing the oocyte with acetate (n = 10, mean ± SD). (C) pH-dependency of relative voltage-dependent open relative probabilities (rel. PO) of MpBK2a-mediated currents prior and after cytosolic acidifications. The increase in the cytoplasmic pH shifted the half-maximal open probability to more negative membrane potentials (n = 6, mean ± SD).

## DISCUSSION

In this study, we present the first characterisation of a plant BK channel. Similar to animal BK channels, MpBK2a is a strongly K^+^-selective, outward-rectifying channel, gated by intra and extracellular pH. MpBK2a requires only trace amounts of external K^+^ for the channel to be open [43]. This is in contrast to Shaker-type Kv channels like *Arabidopsis thaliana* GORK and SKOR, for example, where higher submillimolar external K^+^ concentrations are required to avoid channel conductance collapse [36, 44]. As determined by patch-clamp experiments, the single-channel conductance of MpBK2a (~40 pS outward; ~80 pS inward) is larger than that of typical animal or plant K_v_ channels, yet it is markedly smaller than the conductance of hSlo1 (~200-300 pS) for example [36, 37, 45, 46]. Structural comparison of BK and K_v_ channels revealed that animal BK channel α-subunits, unlike those of K_v_ channels, possess two glutamate residues at the end of the S6 helix (see **Figure S2**). In the functional BK channel tetramer this results in a total of eight negative charges at the cytosolic entrance of the cytosolic BK channel vestibule. Neutralisation of the two glutamate residues by site-directed mutagenesis resulted in a 50% decrease in single-channel conductance when compared to wild-type BK [15 and references therein]. In addition, two negatively charged residues in the outer vestibule affect conductance in hSlo1 [47]. In Marchantia BK channels any of these negatively charged residues are missing and even replaced by positive charges in some cases (**Figure S2**). The lower single-channel conductance of MpBK2a versus animal BK channels may be explained – at least in part – by the lack of a molecuar mechanism whereby negative charges increase K^+^ concentration in the vestibule through long range electrostatic interactions. The lack of these negatively charged glutamate residues additionally would explain our observation that MpBK2a mediates inward currents under certain experimental conditions (see below and **Figure S7**) and possibly explains also why the MpBK2a single-channel outward conductance is smaller than its inward conductance. In hSlo1 the cytosolic glutamate residues stabilise the channel’s closed conformation via ionic intersubunit interactions with positively charged residues of a conserved ‘RKK’ motif [48]. This ‘RKK’ in the TMD-CTD linker region of hSlo1 – upon channel gating – stabilises its open configuration via membrane lipid interaction. The ‘RKK motif is incomplete in the Marchantia BK channel (**Figure S2**) and could explain why V_0.5_ of MpBK2a is about 30 mV compared to ≫ 100 mV for hSlo1. Only with intracellular Ca^2+^, V_0.5_ of hSlo1 is shifted to exhibit a voltage dependence similar to MpBK2a.

Unlike animal BK channels of the Slo1 or Slo2 subfamily, the Marchantia BK channel is neither activated by Ca^2+^, Mg^2+^ nor Na^+^ (**Figure 5 and Figure S6**). Consistent with the results of electrophysiology experiments, structural alignment of the ‘Ca^2+^-bowl region’ revealed that – with the exception of a single conserved glutamine – none of the residues important for Ca^2+^ binding at the RCK2 located ‘Ca^2+^ bowl’ is shared between plant and animal BK channels (**Figure S2**). Likewise, critical residues in the second Ca^2+^-binding site located within the RCK1 domain are essentially absent from the Marchantia BK channels. In animal BKs, coupling between the Ca^2+^ and the voltage sensor is determined by the TMD-CTD linker on one side and the RCK1 N-lobe and the S4–S5 linker on the other [26, 49]. Although the TMD-CTD linker is well conserved between animal and plant BK channels, residues in the RCK1 N-lobe required for either coordination of Ca^2+^ or Mg^2+^ are not. The voltage-dependence of MpBK2a determined in this study, roughly corresponds to the voltage-dependence of hSlo1 under elevated cytosolic Ca^2+^ (see above). Despite the apparent lack of a regulatory mechanism by Ca^2+^_cyt_ in the liverwort BK channel, this may suggest that the prevailing conformation of its CTD structure provides to keep the animal BK channel in its closed state at the hyperpolarised resting membrane potentials typical for plant cells [5], and to respond with fast opening at depolarising membrane potential deflections.

Together with the observation that MpBK2a does not noticeably respond to any of the known hSlo1 inbibitors (IbTx, PenA, Pax) and some prototypic BK openers [25] (**Figure S5**), these findings further distinguish MpBK2a from its animal orthologues. Likewise, the block by extracellular TEA^+^ is rather weak for MpBK2a (~40% block at 10 mM; **Figure S4**) compared to hSlo1 (50% block at 250 μM; [50]), which might be attributed to a missing tyrosine residue replaced by threonine in the outer pore vestibule of the Marchantia BK2a channel (**Figure S2**).

As recently shown for mSlo1 [51], the MpBK2a channel was inhibited by extracellular acidification, similar to its animal counterpart (**Figure 6 and S7**). Acidic residues in the S1-S2 linker (D133A, D147A, D153A) on the extracellular side of each hSlo1 α-subunit have been identified to account for the observed pHO-dependent gating shift (**Figure S2**) and are conserved in other animal voltage-dependent cation channels [51] but not in the MpBK2a (**Figure S2**). However, other acidic residues are present – especially in the S3-S4 linker and the S5-PH loop – and predicted to be extracellularly-accessible that could fulfil a pH sensing role. This suggests independent evolution of pH-dependent gating in plant and animal BK channels. It should be noted that the inhibition of K^+^ efflux by external protons is a phenomenon shared between the Marchantia BK and Shaker-type GORK/SKOR-type K^+^_out_ channels [36, 52]. However, in Arabidopsis GORK/SKOR, the external H^+^ concentration affects the number of channels active rather than their voltage-dependent gating [36]. Opposite to extracellular H^+^, increasing [H^+^]_cyt_ promotes MpBK2a activity by shifting the voltage-dependent gating of MpBK2a toward negative membrane potentials and increasing the open channel probability (**Figure 7**). This H^+^_cyt_-dependent activation of MpBK2a contrasts plant Shaker-type K^+^_out_ channels, known to be strongly inhibited by cytoplasmic acidification [36].

MpBK channels can offer independence from the requirement for more than trace amounts of external potassium for activation, useful in plant cells that are directly exposed to the environment and pH-dependent gating, enabling a means of fine-tuning channel activity synergistically with membrane voltage (cf. refs [10, 11, 36, 53, 54]).

The presence of BK channel genes in streptophyte algae, the closest living algal relatives to land plants, suggest that green BK channels were present in the early plant pioneers that conquered land over 500 Mya, and conserved until late during the divergence of the major land plant groups, before being lost in the angiosperm lineage prior to massive radiation of this plant clade ~200-250 Mya [55]. Future examination of the mutant phenotypes and high-resolution expression data will provide further insight into the roles(s) of BK channels in early plants and during terrestrialisation.

## ACKNOWLEDGMENTS

We thank Kerstin Neuwinger and Katharina Adam for technical assistance. This work was funded in part by DFG grants for the priority programme ‘MAdLand – Molecular Adaptation to Land: Plant Evolution to Change’ to R.H. and D.B., and for the project ‘Evolution of molecular mechanisms that control stomatal closure’ (HE1640/40-1) to R.H.. S.H. acknowledges support by DFG HE2993/18-1. F.S. acknowledges support from an Australian Research Council Discovery Early Career Award (DE200101133) funded by the Australian Government.

## MATERIAL AND METHODS

### Phylogenetic analyses

An HMM profile for Slo channels was created from MAFFT-aligned full-length protein sequences for hSlo2.1 (NP_001274748), mSlo2.1 (NP_001074496), hSlo2.2 (XP_011517180), mSlo2.2 (ACM90117), dSlo2 (NP_001260863), nSlo2 (NP_001257144) and hSlo3 (NP_001027006) using the hmmbuild function of HMMER. BK/Slo-like protein sequences were obtained from the sequence resources detailed in Table S1 using this profile with the hmmsearch function of HMMER and a significance E-value threshold of 1×10^−50^ (sufficient to exclude other animal and plant potassium channel types). A full-length MAFFT alignment of the obtained protein sequences, trimmed using Gblocks (with all options for reduced stringency selected) via the online server at http://molevol.cmima.csic.es/castresana/Gblocks_server.html [56], was used to calculate a maximum likelihood phylogenetic tree using PhyML 3.0 at http://www.atgc-montpellier.fr/phyml/ with SmartModel Selection and 1000 bootstrap replicates [57, 58]. The resulting gene tree was reconciled using TreeFix v1.1.10 [59], using the species tree shown in Figure S1B. Bootstrap values, shown above each branch, were estimated with 100 bootstrapping steps in TreeFix and reflect support for topology and events. Full sequence and species details are given in Table S1.

### 3D modelling of MpBK2a

A 3D homology model of MpBK2a was obtained on the basis of the crystal structure of *Aplysia californica* Slo1 (aSlo1, PDB entry 5TJ6; [26]) using the modelling macro hm_build.mcr of the software package Yasara Structure version 20.12.24 (https://www.yasara.org, [60]). Briefly, the amino acid sequence of MpBK2a covering residues 70-719 and 833-1188 was aligned to the sequence of aSlo1 using a PSI-BLAST search against the RCSB databank with a maximum E-value of 0.5 for template consideration. Five potential templates including entire BK channels or intracellular gating-ring structures from *Homo sapiens* and *Aplysia californica* were identified. For modelling and secondary structure prediction, a target sequence profile was first built against related UniRef90 sequences. Twenty-one initial homology models were then built using six different templates (PDB entries 6V5A, 3MT5, 6V22, 5U70, and 5TJ6) employing up to five slightly different sequence alignments that differed mainly in the adjustments of loop regions. Of those, one exhibited the best overall Z-score of −1.728 in the Yasara scoring routine after energy minimization and molecular dynamics refinement.

### Cloning of the Marchantia *BK* channel gene

The coding sequence (CDS) of *Marchantia polymorpha* BK2a channel (gene details in Table S1) were synthesised by Invitrogen GeneArt Gene Synthesis (ThermoFisher Scientific). For heterologous expression of MpBK2a in *Xenopus* oocytes or HEK cells, the synthesised cDNA was cloned into expression vectors based on pGEM and pcDNA3 vectors, respectively. For cloning purpose, the advanced uracil excision-based cloning technique was used as previously described [61]. Thereby, the CDS was attached to USER-specific 5` and 3` overhangs. PCR conditions were as previously described [62]. The USER-PCR products was treated with the USER enzyme (New England Biolabs, Ipswich, MA, USA) to remove all uracil residue, thus generating single-stranded overlapping ends. Following uracil excision, recirculation of the plasmid was performed at 37 °C for 30 minutes followed by 30 minutes at room temperature. Constructs were then immediately transformed into chemical competent *Escherichia coli* cells (XL1-Blue MRF’). All constructs were verified by sequencing. For functional analysis in *Xenopus* oocytes, cRNAs were prepared using the AmpliCap-Max T7 High Yield Message Maker Kit (Cellscript, Madison, WI, USA) by following the manufacturer’s instructions and 10 ng of cRNA was injected into selected oocytes. Oocytes were incubated for 2 days at 16 °C in ND96 (10 mM HEPES pH 7.4, 96 mM NaCl, 2 mM KCl, 1 mM MgCl_2_, 1 mM CaCl_2_) solution containing gentamycin.

### *Xenopus* oocyte preparation

Investigations on MpBK2a channel properties were performed in oocytes of the African clawfrog *Xenopus laevis*. Permission for keeping *Xenopus* exists at the Julius-von-Sachs Institute and is registered at the government of Lower Franconia (Reference Number 55.2-2532-2-1035). For oocyte isolation, mature female *laevis* frogs were anesthetised by immersion in water containing 0.1% 3-aminobenzoic acid ethyl ester. Following partial ovariectomy, stage V or VI oocytes were treated with 0.14 mg/ml collagenase I in Ca^2+^-free ND96 buffer (10 mM HEPES pH 7.4, 96 mM NaCl, 2 mM KCl, 1 mM MgCl_2_,) for 1.5 h. Subsequently, oocytes were washed with Ca^2+^-free ND96 buffer and kept at 16 °C in ND96 solution (10 mM HEPES pH 7.4, 96 mM NaCl, 2 mM KCl, 1 mM MgCl_2_, 1 mM CaCl_2_) containing 50 mg/l gentamycin [8].

### Two Electrode Voltage Clamp technique on MpBK2a-expressing oocytes

In Two Electrode Voltage Clamp (TEVC) studies, MpBK2a-expressing oocytes were perfused with standard bath solution containing 10 mM MES/Tris (pH ≥ 5.6), 1 mM Ca^2+^, 1 mM Mg^2+^ and 1 mM LaCl_3_ and 100 mM K^+^ (Na^+^ or Li^+^ for selectivity analysis). The ionic strength for measurements under varying potassium concentrations was balanced with lithium to give a total concentration of 100 mM. The corresponding salts were either gluconate- or chloride-based and the blocking substances were also used as chloride salts and added in concentrations as indicated in the figures. For cytosolic acidification, oocytes were perfused with bath solutions containing 10 mM K^+^ in the presence of 30 mM NaOAc. The osmolality of each solution was adjusted to 220 mOsm/kg using D-sorbitol. The standard pulse protocol was applied in 20 mV increments from −120 to 60 mV starting from a holding potential (V_H_) of −100 mV. Any deviations of this standard pulse protocol are indicated and mentioned in the figure and legends. The steady-state currents were extracted at the end of the test pulses and used for current-voltage relations (IV). Currents were normalised to conditions as described in the figure. For analysing the reversal potential (E_rev_) tail currents (I_tail_) at the beginning of the test pulses after a pre-activation of 40 mV were recorded after the capacitive currents. To estimate the relative open probability, I_tail_ were extracted immediately after the voltage jump to 50 mV from the applied test pulses. For calculation of the relative open probability, data points were fitted with a Boltzmann function resulting in V_0.5_, the half-maximal open probability [63, 64]. To record the current during the cytosolic acidification the following voltage ramp was used: starting from V_H_ at −100 mV to a test pulse at 40 mV, the voltage ramp ranged from 40 to −140 mV for 100 ms followed by a post-pulse at −140 mV before going back to the V_H_. For statistical analysis and graph preparation, the software Igor Pro (WaveMetrics, Lake Oswego, OR) was used. The results are presented as mean ± standard deviation to measure the dispersion of a dataset relative to its mean (SD). n is the number of independent measurements.

### Patch-clamp experiments on *Xenopus* oocytes

Oocytes were injected with MpBK2a cRNA 3-4 days prior to patch-clamp experiments. Patch pipettes were pulled from borosilicate glass tubing (outer diameter 2.0 mm, inner diameter 1.0 mm, resistance 0.7-1.2 MΩ). For sealing, the external buffer contained 100 mM KCl, 10 mM HEPES, 1 mM EGTA at pH 7.4, whereas patch-clamp measurements were performed in either potassium-free (100 mM NaCl, 1 mM EGTA, 10 mM HEPES, pH 7.4 (NaOH)) or potassium-based buffers (100 mM KCl, 1 mM EGTA, 10 mM HEPES, pH 7.4 (KOH)). The pipette solution contained 100 mM KCl, 1 mM CaCl_2_, and 10 mM MES, pH 5.6 (KOH). After obtaining the inside-out patches, the pipette was positioned in front of a multibarrel application system that was used to provide different solutions to the cytosolic face of the patch. Ion currents were recorded with the patch-clamp technique using an EPC 10 USB patch-clamp amplifier (HEKA Elektronik, Lambrecht, Germany). Current measurements were controlled, and data were collected with the PatchMaster software V2×90 (HEKA Elektronik). The voltage protocols for recording current amplitudes was as follows: (i) For steady-state current recordings: starting from a holding potential of −60 mV single 500 ms voltage pulses from −60 mV to 80 mV in 10 mV steps were applied, followed by a 250 ms voltage pulse at 80 mV and a 2 s voltage pulse at −60 mV. Steady-state currents were taken at the end of the respective voltage pulse. (ii) For recording tail currents: starting from a holding potential of −60 mV channels were preactivated by a 2 s lasting voltage pulse at 80 mV and tail currents were taken directly after a voltage jump to 2 s lasting single voltage pulses from 80 mV to −60 mV in 10 mV decrements. The sampling rate was 50 kHz and a 2.9 kHz Bessel filter was used.

### Patch-clamp experiments on HEK293T cells

Marchantia polymorpha BK2a (Mp7g12080) was transiently expressed from a vector with CMV promoter in human embryonic kidney (HEK) 293T cells (CAMR), cultured in Dulbecco’s modified Eagle’s medium containing 45% Ham’s F12 medium (PAA) and 10% fetal bovine serum in a 5% CO_2_ incubator at 37 °C. Cells were trypsinised, diluted with culture medium, and seeded on 12-mm glass coverslips 1 day prior to transfection. Cells were transfected with the respective plasmids using the Rotifect (Roth) transfection reagent. Co-transfection of a plasmid coding for sfGFP/spGFP was used for identification of transfected cells by means GFP fluorescence. After transfection, cells were maintained in an incubator at 26 °C. Electrophysiological experiments were performed 2−6 days after transfection at room temperature (20-22 °C).

Patch pipettes were fabricated from borosilicate class, coated with dental wax to reduce capacitance, and polished to yield resistances of 1-2 MΩ. Electrophysiological recordings were performed with an EPC10 amplifier, operated by PatchMaster software (HEKA Elektronik). In whole-cell recordings, the series resistance was compensated electronically up to 85%. Electrophysiological data were analysed with FitMaster (HEKA Elektronik) and IgorPro (WaveMetrics).

For the recording of macroscopic currents in the whole-cell mode, a p/n protocol for linear leak and capacitive current correction was applied. Voltage pulse protocols and solutions are indicated in the respective figures. Voltage-dependence of channel activation was described with a Boltzmann-type function:

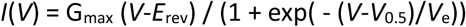

where G_max_ is the maximal conductance, *E*_rev_ the reversal potential, *V*_0.5_ the half-activation voltage, and *V*_e_ the slope factor indicating voltage dependence. For tail current-voltage relationships, G_max_ (*V*-*E*_rev_) was replaced by the normalized maximal tail current. Activation and tail current kinetics was estimated with single-exponential functions.

Single-channel open probability (P_O_) was estimated using all-points amplitude histograms constructed from the data traces, manually corrected for leak current. Single-channel conductance was estimated by linear fits to *i*(*V*) relationships, separately in the positive and negative current direction.

Iberiotoxin was dissolved in bath solution (100 μM), all other Slo1 channel modulators were dissolved in DMSO (1-10 mM). Typical Slo1 channel inhibitors and activators, freshly diluted before the experiment, were applied by full bath exchange when recording currents in the whole-cell mode. NS-compounds were from TOCRIS Bioscience, all other modulators were from Sigma-Aldrich.

The results are presented as mean ± standard error of the mean (SEM) (n) where n is the number of independent measurements.

### Localisation studies of MpBK2a by transient transformation of *N. benthamiana*

To generate the MpBK2a::YFP construct, the full-length USER-PCR (described in `Cloning of the Marchantia *BK* channel gene’) was cloned into the binary vector pCAMBIA 2300 (35S:USER::YFP:35STerm) using the advanced uracil excision-based cloning technique [61, 62]. *Agrobacterium tumefaciens* cells of strain GV3101 were used for transformation and the preparation of the bacterial suspension is described in [65]. For transient transformation, *Nicotiana benthamiana* leaves of 3-week-old plants were infiltrated at their abaxial side with the bacterial suspension. Because of the co-infiltration of two bacterial suspensions carrying the MpBK2a::YFP constructs and the plasma membrane marker, a ratio of 2:1 was used. As plasma membrane marker the DsRed2-labelled REMORIN 1.3 from Arabidopsis (DsRed2::AtREM1.3) [34] was used. The transient transformed plants were incubated for 3 days in a growth chamber before microscopy was performed. A confocal laser scanning microscope (TCS SP5 2; Leica) and the water immersion objective lens Leica HCX IRAPO L25×/0.95W were used to visualise YFP fluorescence (excitation: 488nm; emission: 514–551nm) and DsRED2 fluorescence (excitation: 561nm, emission: 560-600nm). Images were taken using the software LAS AF (Leica Application Suite Advanced Fluorescence 2.4.1; Leica).

