## Supplementary material for "The liverwort *Marchantia polymorpha* operates a depolarization-activated Slowpoke (SLO) K^+^ channel that recognises pH changes in the environment": suppl. figures and tables

**Supplementary Information:**

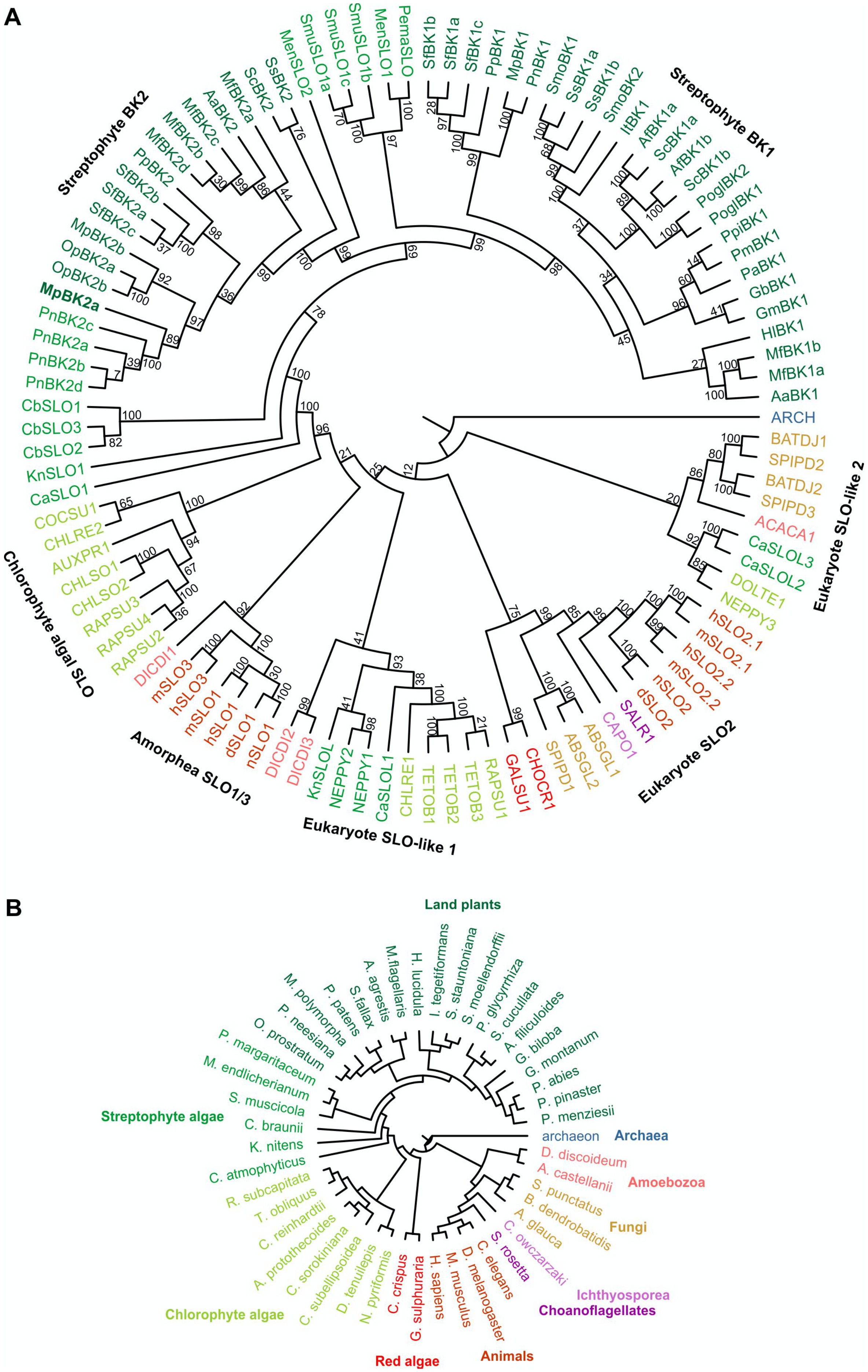
A
Streptophyte BK2
Streptophyte BK1
Chlorophyte algal SLO
Amorphea SLO1/3
Eukaryote SLO-like 1
Eukaryote SLO2
Eukaryote SLO-like 2
MenSLO2
SmuSLO1a
SmuSLO1c
SmuSLO1b
MenSLO1
PemaSLO
SfBK1b
SfBK1a
SfBK1c
PpBK1
MpBK1
PnBK1
SmoBK1
SsBK1a
SsBK1b
SmoBK2
ItBK1
AfBK1a
ScBK1a
AfBK1b
ScBK1b
PoglBK2
PoglBK1
PpiBK1
PmBK1
PaBK1
GbBK1
GmBK1
HlBK1
MfBK1b
MfBK1a
AaBK1
ARCH
BATDJ1
SPIPD2
BATDJ2
SPIPD3
ACACA1
CaSLOL3
CaSLOL2
DOLTE1
NEPPY3
hSLO2.1
mSLO2.1
hSLO2.2
mSLO2.2
nSLO2
dSLO2
SALR1
CAPO1
ABSGL1
ABSGL2
SPIPD1
CHOCR1
GALSU1
RAPSU1
TETOB3
TETOB2
TETOB1
CHLRE1
CaSLOL1
NEPPY1
NEPPY2
KnSLOL
DICDI3
DICDI2
nSLO1
dSLO1
hSLO1
mSLO1
hSLO3
mSLO3
DICDI1
RAPSU2
RAPSU4
RAPSU3
CHLSO2
CHLSO1
AUXPR1
CHLRE2
COCSU1
CaSLO1
KnSLO1
CbSLO2
CbSLO3
CbSLO1
PnBK2d
PnBK2b
PnBK2a
PnBK2c
MpBK2a
OpBK2b
OpBK2a
MpBK2b
SfBK2c
SfBK2a
SfBK2b
PpBK2
MfBK2d
MfBK2b
MfBK2c
AaBK2
MfBK2a
ScBK2
SsBK2
B
Land plants
Streptophyte algae
Chlorophyte algae
Red algae
Animals
Choanoflagellates
Ichthyosporea
Fungi
Amoebozoa
Archaea
M. polymorpha
P. patens
S. fallax
A. agrestis
M. flagellaris
H. lucidula
I. tegetiformans
S. stauntoniana
S. moellendorffii
P. glycyrrhiza
S. cucullata
A. filiculoides
G. biloba
G. montanum
P. abies
P. pinaster
P. menziesii
archaeon
D. discoideum
A. castellanii
S. punctatus
B. dendrobatidis
A. glauca
C. owczarzaki
S. rosetta
C. elegans
D. melanogaster
M. musculus
H. sapiens
G. sulphuraria
C. crispus
N. pyriformis
D. tenuilepis
C. subellipsoidea
C. sorokiniana
A. protothecoides
C. reinhardtii
T. obliquus
R. subcapitata
C. atmophyticus
K. nitens
C. braunii
S. muscicola
M. endlicherianum
P. margaritaceum
O. prostratum
P. neesiana

**Figure S1. Two BK subfamilies are encoded in land plants including *Marchantia polymorpha*.**

(A) The fully-expanded phylogeny of the eukaryote Slo family shown in Figure 1A, rooted to a related archaeon gene. Bootstrap values, shown above each branch, reflect support for topology and events, as estimated with 100 bootstrapping steps in TreeFix [1]. Full gene details are given in Table S1. (B) The species tree used for gene tree reconciliation.

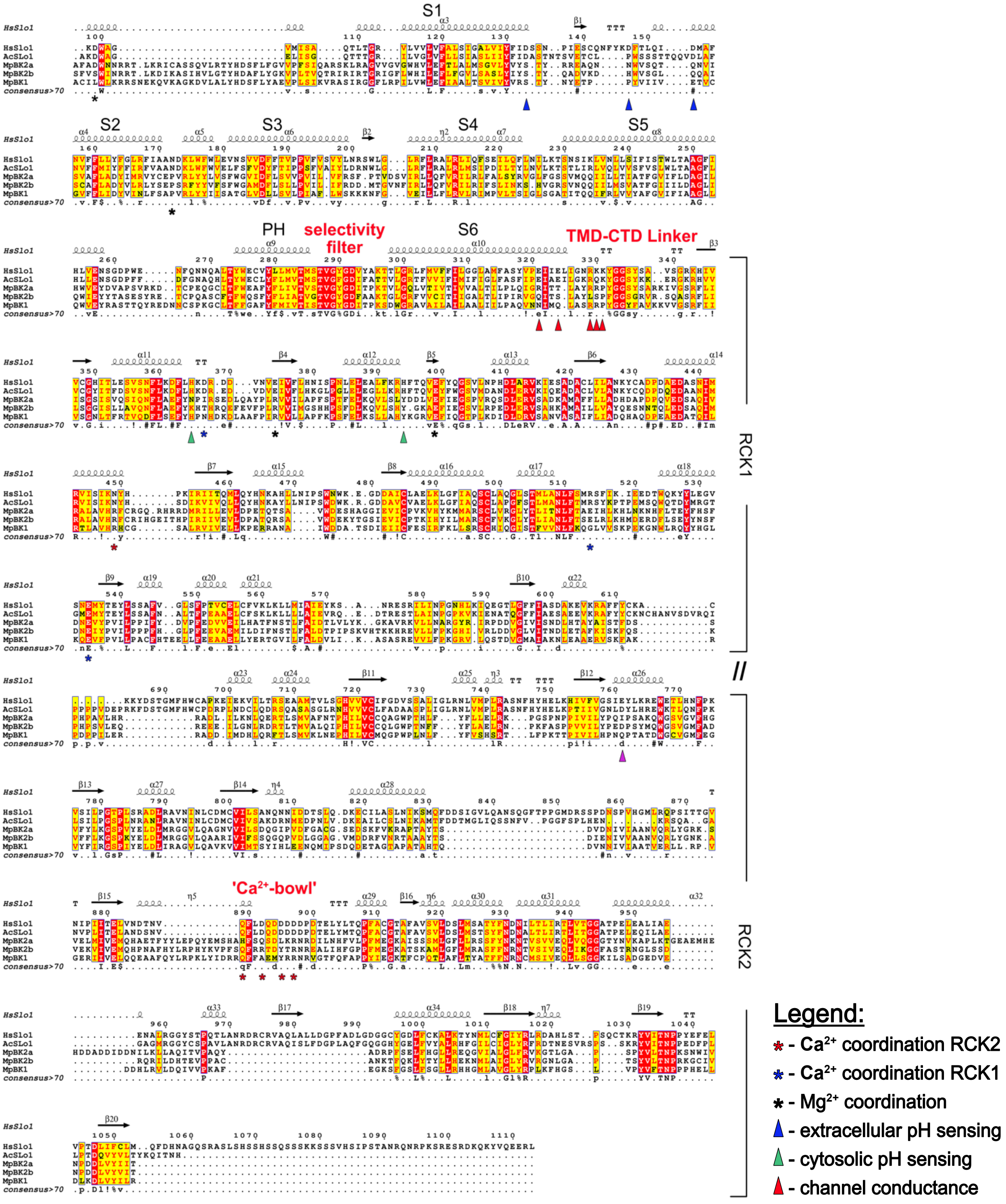


**Figure S2: Full-length, structurally-guided alignment of Marchantia and animal BK channel α–subunits**

Amino acid sequences of hSlo1, AcSlo1, MpBK2a, MpBK2b, and MpBK1 were aligned using the Expresso mode of T-Coffee [2]. ESPript (v3.0) [3] was employed for secondary structure prediction and annotation of the alignment guided by the hSlo1 structure (PDB entry 6V38). Relative positions of the transmembrane domains

(S1 – S6), the pore helix (PH) and the RCK domains are indicated. Functional residues are denoted as in the legend.

On the basis of site-directed mutagenesis and structural studies, two distinct high-affinity $Ca^{2+}$-binding sites have been identified in the CTD of animal Slo1 α subunits [4-8]. The first site, also known as the '$Ca^{2+}$ bowl site' is located near the interface between RCK domains of adjacent subunits. Here, within the RCK2 domain, the side chains of two aspartic residues together with the main-chain carbonyl oxygen atoms of neighbouring glutamine and aspartic residues as well as the side chain of aspartic acid from the RCK1 domain of the neighbouring Slo1 α–subunit are essential for $Ca^{2+}$ coordination. Structural data further show that the $Ca^{2+}$-sensory CTD moiety is directly linked to the pore through the S6 linker region and indirectly through the non-covalent interfaces formed between the RCK1 N-lobe and the voltage sensor/S4–S5 linker [4, 6] and that this well-formed interface provides for coupling between the $Ca^{2+}$ sensor and the voltage sensor.

In addition to micromolar $Ca^{2+}_{cyt}$, millimolar intracellular $Mg^{2+}$ activates animal Slo1 channels, while Slo2 channels are activated by high intracellular $Na^{+}$ concentrations [9]. $Mg^{2+}$ binding involves residues in the VSD and the CTD from two neighbouring subunits forming an interdomain binding site [4] and $Na^{+}$ binding was shown to depend on a single aspartate residue in the CTD of hSlo2.1 [10]. Consistent with the structural model and the lack of sequence conservation, our electrophysiological analyses demonstrated that the Marchantia BK2a is neither affected by cytosolic, millimolar concentrations of $Mg^{2+}$ nor $Na^{+}$.

Two conserved histidine residues in the RCK1 domain of hSlo1 have been identified to communicate changes in $pH_{cyt}$ between the CTS and VSD and thus affect voltage-dependent gating of the animal BK channel [11]. None of these histidines is conserved between hSlo1 and MpBK2a channels, though several further histidines in the RCK1 domain of the Marchantia BK channel may provide to $H^{+}_{cyt}$ susceptibility.

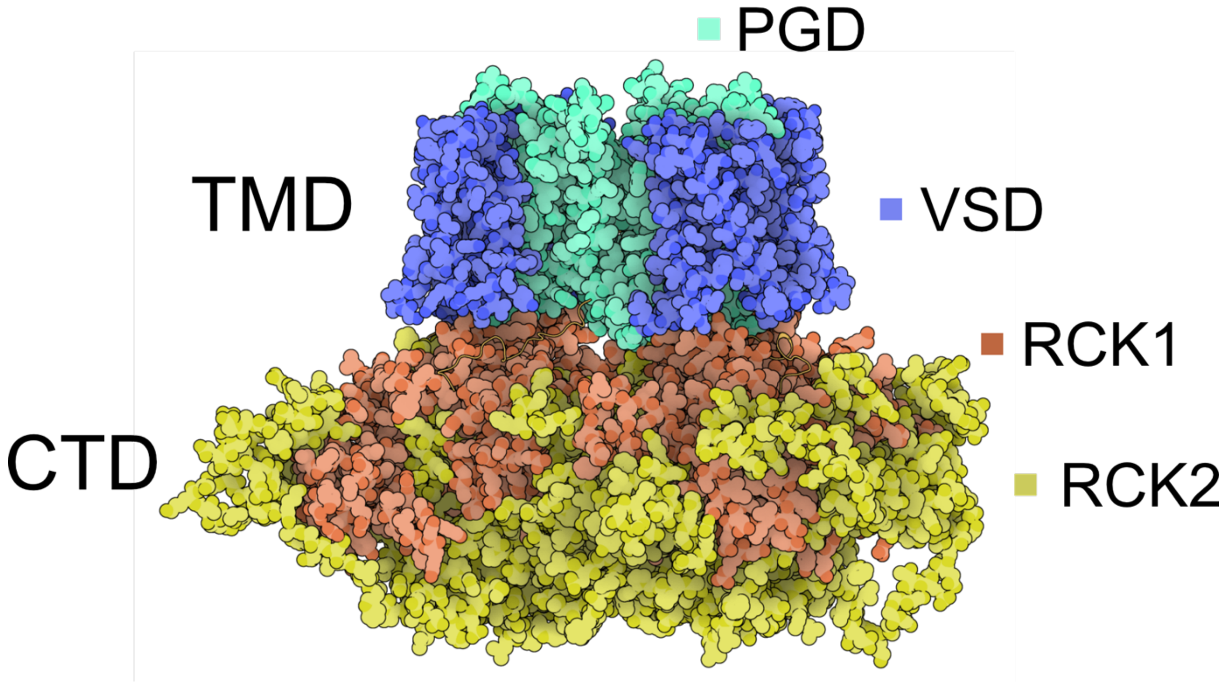


**Figure S3. Structural model of MpBK2a based on Aplysia Slo1**

Cartoon of the MpBK2a channel based on structural modelling (see methods) and generated with The Protein Imager (https://3dproteinimaging.com/; [12]). The voltage-sensing domain (VSD; blue) and the pore gate domain (PGD; green) together constitute the transmembrane domain (TMD). The cytosolic tail domain (CTD) is composed of the RCK1 (red) and RCK2 (yellow) domains.

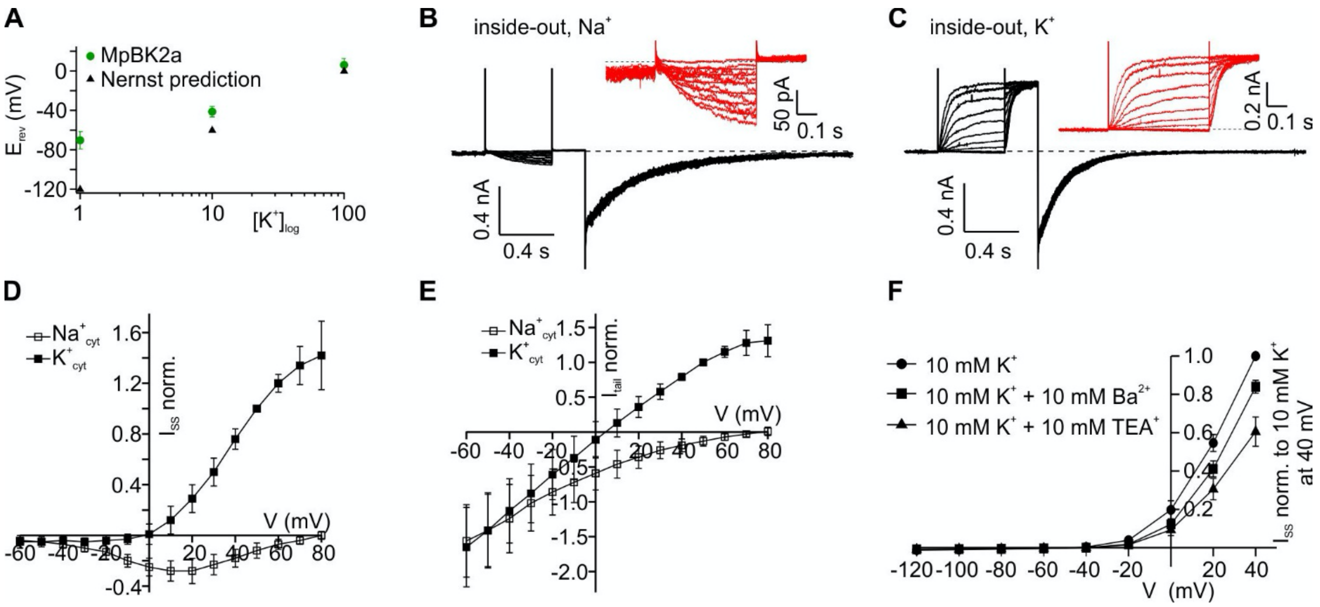


**Figure S4. MpBK2a is $K^+$ selective and is blocked by $TEA^+$ and $Ba^{2+}$.**

(A) The reversal potentials ($E_{rev}$) mediated by MpBK2a (green) were obtained from $I_{tail}$ recordings in chloride-based solutions at pH 5.6 after a pre-activation at 40 mV. The resulting $E_{rev}$ plotted against the external $K^+$ concentration (logarithmic scale) shifted in $K^+$ dependent manner comparable to the predicted equilibrium potential for $K^+$ according to the Nernst equation (n = 5, mean ± SD). (B-E) Cation selectivity was investigated by patch-clamp analysis using *Xenopus* oocytes expressing MpBK2a. Measurements were performed in the inside-out configuration with 100 mM potassium in the patch pipette. The cytosolic face of the oocyte membrane patch was perfused with either (B) sodium- or (C) potassium-based buffers. Starting from a holding potential of -60 mV, single 500 ms voltage pulses ranging from -60 mV to 80 mV were applied in 10 mV steps followed by a 250 ms voltage pulse at 80 mV and a 2 s voltage pulse at -60 mV. In the absence of potassium in the bath (B), representing the cytosol, only inward currents could be recorded, indicating potassium flux from the pipette into the bath. When perfusing potassium-based buffers (C), macroscopic outward currents appeared at positive voltages, representing potassium flux from the bath into the pipette. Representative current traces are shown. (D) Steady-state currents from measurements shown in (A) and (B) plotted against the applied voltage ($n \geq 4$, mean ± SD). (E) Tail currents at voltages ranging from -60 mV to 80 mV recorded directly after a 2 s lasting 80 mV pre-activation pulse are plotted against the applied voltage. Inside-out membrane patches of oocytes expressing MpBK2a were perfused with either sodium- or potassium-based buffers ($n \geq 4$, mean ± SD). (F) Steady-state currents ($I_{SS}$) mediated by MpBK2a in a voltage range from -120 to 40 mV and in the presence of 10 mM $K^+_{ext}$ and the indicated blocker (10 mM each). The application of $TEA^+$ and $Ba^{2+}$ reduced the current by about 40% and 20%, respectively ($n \geq 8$, mean ± SD).

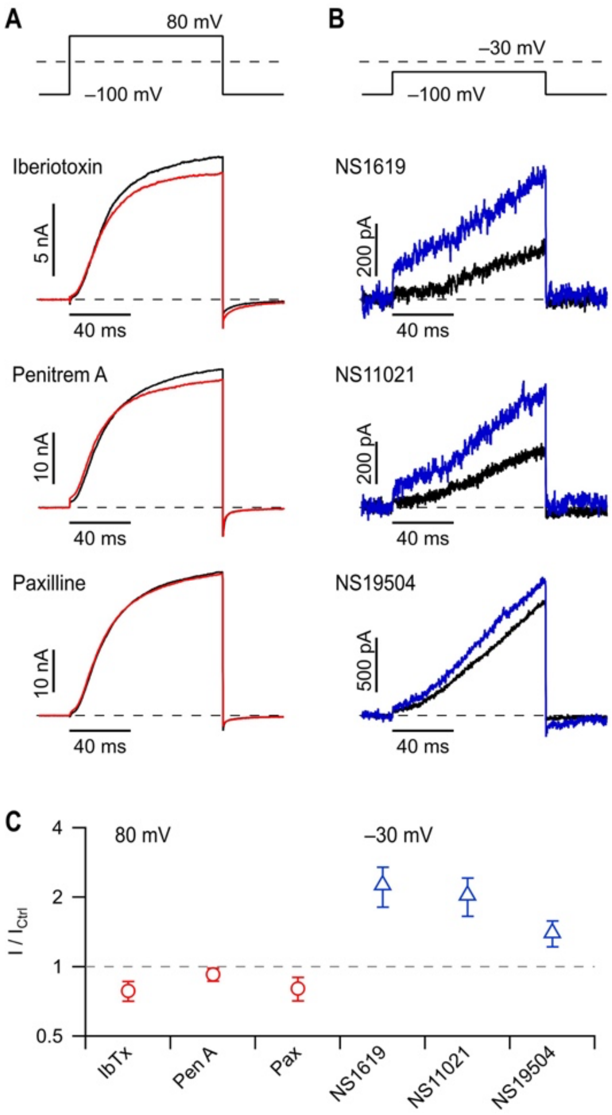


**Figure S5. Typical modulators of human Slo1 BK channels do not noticeably affect MpBK2a.**

(A) Pulse protocol and superposition of whole-cell current traces of MpBK2a before (black) and after (red) application of Slo1 inhibitors iberiotoxin, penitrem A, or paxilline (1 µM each). (B) Current traces before (black) and after (blue) application of Slo1 activators NS1619, NS11021, and NS19504 (10 µM each). (C) Mean relative current after application of Slo1 BK inhibitors (red, at 80 mV) or activators (blue, at - 30 mV). Data points are means ± SEM (n = 5-6).

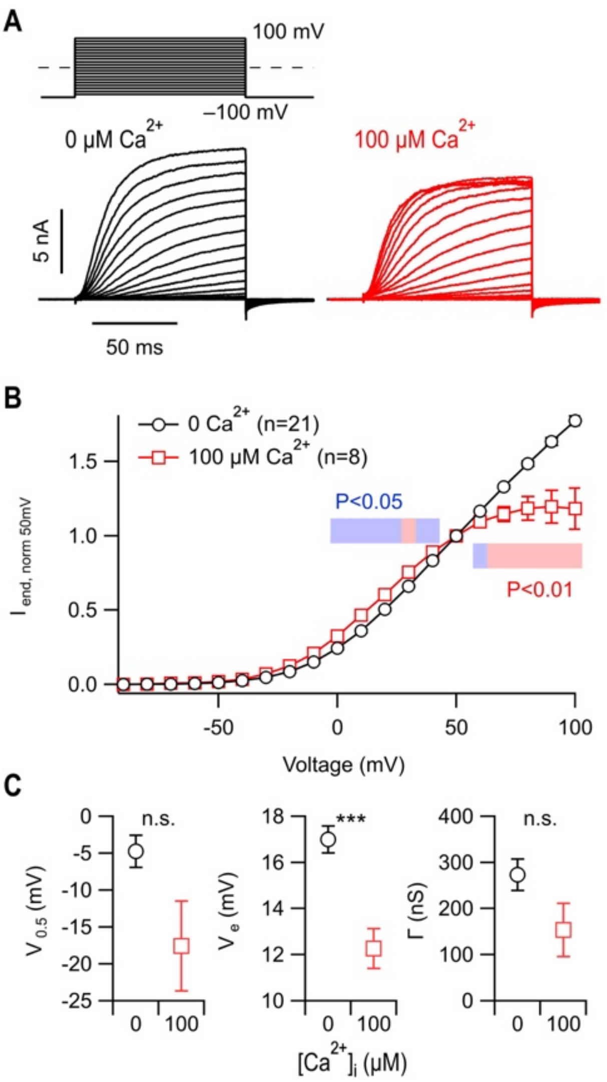


**Figure S6. Marginal impact of intracellular $Ca^{2+}$ on MpBK2a channel gating.**

(A) Superposition of whole-cell current trace from HEK29T cells expressing MpBK2a according to the indicated voltage protocol. Intracellular (pipette) solutions either contained no free $Ca^{2+}$ (left, buffered with 10 mM EGTA) or 100 µM (right). (B) Maximal currents obtained from pulse protocols shown in (A) normalised to the amplitudes obtained at 50 mV without and with intracellular $Ca^{2+}$. Data are means ± SEM (n in parentheses); straight lines connect data points. (C) Analysis parameters characterising the voltage-dependence of channel gating under the conditions of (A) and (B), from left to right: $V_{0.5}$, the half-maximal activation voltage; $V_e$, the slope factor, and $\Gamma$, the conductance.

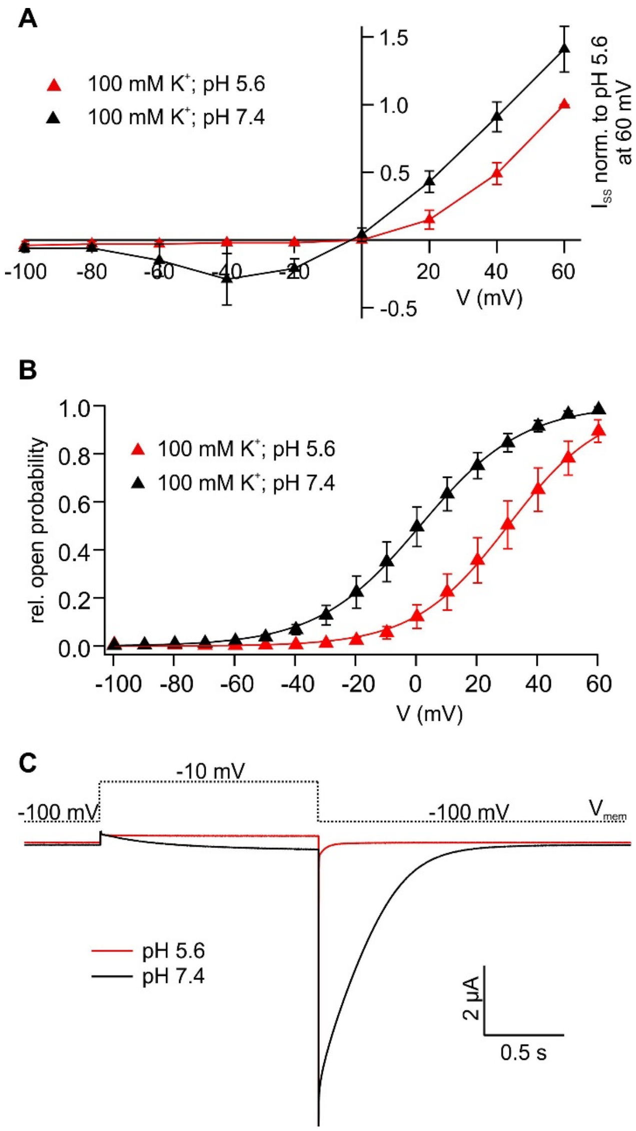


**Figure S7. Extracellular pH modulates MpBK2a current amplitude and kinetics**

(A) $I_{SS}$ recorded in 100 mM $K^+$ at the indicated pH in a voltage range from -100 to 60 mV. In solutions with basic external pH (pH 7.4, black), MpBK2a channels open at more negative membrane potentials, resulting in potassium influx, whereas channels are closed at negative membrane potentials in acidic pH (pH 5.6, red) (n = 5, mean ± SD). (B) The relative open probability of MpBK2a shifts to more negative membrane potentials with increased extracellular pH. Relative open probability was determined from $I_{tail}$ following the test pulses in the voltage range -100 to 60°mV to post-pulse and oocytes were perfused with 100 mM $K^+$ at the indicated pH values. Data points were fitted with a Boltzmann function (n = 5, mean ± SEM). (C) Representative current traces from mediated by MpBK2a in 100 mM $K^+$ solutions at the indicated pH. The channel closure is affected by a change in the extracellular $H^+$ concentration resulting in τ = 0.07 ± 0.02 s in pH 5.6 (n= 5, mean ± SD) and τ = 0.46 ± 0.27 s in pH 7.4 (n = 4, mean ± SD). The dotted line indicates the applied voltages.

**Table S1: Sequence details**.

Sequences were obtained from CNGB (https://db.cngb.org/), ConGenIE (http://congenie.org/), DRYAD (https://datadryad.org/stash), EnsemblPlants (https://plants.ensembl.org/), FernBase (https://www.fernbase.org/), figshare (https://figshare.com/), GenBank (https://www.ncbi.nlm.nih.gov/), GigaDB (http://gigadb.org/), Gymno PLAZA (https://bioinformatics.psb.ugent.be/plaza/versions/gymno-plaza/), JGI (https://genome.jgi.doe.gov/portal/), the *K. nitens* NIES-2285 genome project (http://www.plantmorphogenesis.bio.titech.ac.jp/~algae_genome_project/klebsormidium/index.html), OrcAE (https://bioinformatics.psb.ugent.be/orcae/), PineRefSeq (https://pinerefseq.faculty.ucdavis.edu/), The *Penium* genome database (http://bioinfo.bti.cornell.edu/Penium), UniProt (https://www.uniprot.org/) or as otherwise indicated. The Marchantia BK channel featured in electrophysiology experiments is highlighted in green.

| Group | Species | Gene name | Accession | Sequence sourc | Gene reference |
|---|---|---|---|---|---|
| chaea | archaeon (unclassifie archaea) | ARCH | A0A482SXX1_9ARCH | UniProt | This study |
| noebozoa | ictyostelium discoideu (Slime mold) | DICDI1 | Q55CU6_DICDI | UniProt | This study |
| | | DICDI2 | Q86I16_DICDI | | |
| | | DICDI3 | Q54SN1_DICDI | | |
| | *canthamoeba castella* str. Neff | ACACA1 | L8HDA6_ACACA | | |
| ngi | *Absidia glauca* | ABSGL1 | A0A168QBS7_ABSGL | UniProt | This study |
| | | ABSGL2 | A0A168P632_ABSGL | | |
| | *Batrachochytrium dendrobatidis* (strain JAM81 / FGSC 10211 | BATDJ1 | F4NZW7_BATDJ | | |
| | | BATDJ2 | F4NZ65_BATDJ | | |
| | *pizellomyces punctat* (strain DAOM BR117 | SPIPD1 | A0A0L0HFL3_SPIPD | | |
| | | SPIPD2 | A0A0L0HS60_SPIPD | | |
| | | SPIPD3 | A0A0L0HT08_SPIPD | | |
| thyosporea | *Capsaspora owczarza* (strain ATCC 30864) | CAPO1 | A0A0D2UAP2_CAPO3 | UniProt | This study |
| oanoflagellates | *Salpingoeca rosetta* (strain ATCC 50818 / BSB-021) | SALR1 | F2UHN6_SALR5 | UniProt | This study |
| imals | *Caenorhabditis elegan* | nSlo1 | SLO1_CAEEL | UniProt | /ei, et al. [13]; Yuan, al. [14] |
| | | nSlo2 | H9G2R4_CAEEL | | |
| | *rosophila melanogast* (common fruit fly) | dSlo1 | SLO_DROME | | Elkins, et al. [15]; Atkinson, et al. [16]; |
| | | dSlo2 | A8DY93_DROME | | |
| | *Mus musculus* (mouse | mSlo1 | KCMA1_MOUSE | | Butler, et al. [17]; Schreiber, et al. [18]; Yuan, et al. [19] |
| | | mSlo2.1 | D3Z649_MOUSE | | |
| | | mSlo2.2 | KCNT1_MOUSE | | |
| | | mSlo3 | KCNU1_MOUSE | | |
| | *Homo sapiens* (humar | hSlo1 | KCMA1_HUMAN | | seng-Crank, et al. [2( Pallanck and Ganetzk 1]; Bhattacharjee, et [22] |
| | | hSlo2.1 | KCNT2_HUMAN | | |
| | | hSlo2.2 | KCNT1_HUMAN | | |
| | | hSlo3 | KCNU1_HUMAN | | |
| odophyta (red algae) | *Galdieria sulphuraria* | GALSU1 | M2X6F1_GALSU | UniProt | This study |
| | *Chondrus crispus* | CHOCR1 | R7QF10_CHOCR | | |
| nts: chlorophyte alga | *Auxenochlorella protothecoides* | AUXPR1 | A0A087SHU6_AUXPR | UniProt | This study |
| | *Chlamydomonas reinhardtii* | CHLRE1 | Cre02.g146300 | 5.6 [23], Phytozor v12 [24] | This study |
| | | CHLRE2 | Cre13.g603750 | | |
| | *Chlorella sorokiniana* | CHLSO1 | A0A2P6TKU3_CHLSO | UniProt | This study |
| | | CHLSO2 | A0A2P6TKS3_CHLSO | | |
| | *occomyxa subellipsoic (strain C-169)* | COCSU1 | Csu59753 | 2.0 [25], Phytozon v12 [24] | This study |
| | *olichomastix_tenuilep* | DOLTE1 | scaffold-XOAL-2001448 | kp [26], GigaDB [2 | This study |
| | *lephroselmis pyriform* | NEPPY1 | scaffold-ISIM-2000651 | kp [26], GigaDB [2 | This study |
| | | NEPPY2 | scaffold-ISIM-2035837 | | |
| | | NEPPY3 | scaffold-ISIM-2036937 | | |
| | *Tetradesmus obliquu* | TETOB1 | A0A383V8D8_TETOB | UniProt | This study |
| | | TETOB2 | A0A383W8M2_TETOB | | |
| | | TETOB3 | A0A383W5Q4_TETOB | | |
| | *aphidocelis subcapita* | RAPSU1 | A0A2V0NW56_9CHLO | UniProt | This study |

| Group | Species | Gene name | Accession | Sequence sourc | Gene reference |
|---|---|---|---|---|---|
| | | RAPSU2 | A0A2P6VA34_9CHLO | | |
| | | RAPSU3 | A0A2P6VA49_9CHLO | | |
| | | RAPSU4 | A0A2P6VA38_9CHLO | | |
| ants: streptophyte alga | *hlorokybus atmophytic* | CaSLO1 | Chrsp29S00338 | CNGB assembly CNA0002353 [28 | Wang, et al. [28]; Thi study |
| | | CaSLOL1 | Chrsp12S01991 | | |
| | | CaSLOL2 | Chrsp1S03195 | | |
| | | CaSLOL3 | Chrsp36S00420 | | |
| | *Mesostigma viride* | - | - | CNGB assembly CNA0002352 [28 | - |
| | *Klebsormidium nitens* | KnSLO1 | kfl00088_0200_v1.1 | *. nitens* NIES-228 enome project v1 [29] | This study |
| | | KnSLOL | kfl00409_0060_v1.1 | | |
| | *Chara braunii* | CbSLO1 | g30987 | v20170414, OrcA [30] | Nishiyama, et al. [30] Sussmilch, et al. [31] |
| | | CbSLO2 | g32413 (NT) + g32414 (CT | | |
| | | CbSLO3 | g32402 | | |
| | *Mesotaenium endlicherianum* | MenSLO1 | ME000198S02739 | Figshare [32] | This study |
| | | MenSLO2 | ME000148S01122 | | |
| | *Penium margaritaceu* | PemaSLO | pm010361.t1 SLO | he *Penium* genon database [33] | |
| | *Spirogloea muscicola* | SmuSLO1a | SM000146S00940 | Figshare [32] | |
| | | SmuSLO1b | SM000072S21206 | | |
| | | SmuSLO1c | SM000048S16560 | | |
| ants: bryophytes: erworts | *Marchantia polymorph* | MpBK1 | OAE35698 (improved annotation of Mapoly0007s0089) | GenBank, 3.1 [34] Phytozon [24] | Sussmilch, et al. [31] |
| | | **MpBK2a*** | **Mapoly0003s0221** | 3.1 [34], Phytozon [24] | |
| | | MpBK2b | Mapoly0159s0025 | | |
| | *dontoschisma_prostra m* | OpBK2a | scaffold-YBQN-2010875 | kp [26], GigaDB [2 | This study |
| | | OpBK2b | scaffold-YBQN-2010876 | | |
| | *Pellia_neesiana* | PnBK1 | scaffold-JHFI-2014379 | kp [26], GigaDB [2 | This study |
| | | PnBK2a | scaffold-JHFI-2009591 | | |
| | | PnBK2b | scaffold-JHFI-2009592 | | |
| | | PnBK2c | scaffold-JHFI-2009593 | | |
| | | PnBK2d | scaffold-JHFI-2009595 | | |
| ants: bryophytes: osses | *Sphagnum fallax* | SfBK1a | Sphfalx02G147800.1.p | *phagnum fallax* v1 OE-JGI, Phytozon v12 [24] | This study |
| | | SfBK1b | Sphfalx05G064000.1.p | | |
| | | SfBK1c | Sphfalx01G177000.1.p | | |
| | | SfBK2a | Sphfalx03G074000.1.p | | |
| | | SfBK2b | Sphfalx14G038400.1.p | | |
| | | SfBK2c | Sphfalx13G076100.1.p | | |
| | *Physcomitrella paten* | PpBK1 | Pp3c8_8470V3.1 | Phypa v3 [35], EnsemblPlants | mez-Porras, et al. [3 Sussmilch, et al. [31] |
| | | PpBK2 | Pp3c27_7700V3.1 | | |
| ants: bryophytes: rnworts | *Anthoceros agrestis* | AaBK1 | agrBONN_evm.model.Sc2 wM_362.1151.1 | rnworts.uzh.ch/en nwort-genomes.ht 7] | This study |
| | | AaBK2 | agrBONN_evm.model.Sc2 wM_228.1525.1 | | |
| | *Megaceros flagellaris* | MfBK1a | scaffold-UCRN-2003202 | kp [26], GigaDB [2 | This study |
| | | MfBK1b | scaffold-UCRN-2003201 | | |
| | | MfBK2a | scaffold-UCRN-2012193 | | |
| | | MfBK2b | scaffold-UCRN-2005362 | | |
| | | MfBK2c | scaffold-UCRN-2005360 | | |
| | | MfBK2d | scaffold-UCRN-2005361 | | |
| ants: lycophytes | *Huperzia lucidula* | HlBK1 | scaffold-GKAG-2022102 | kp [26], GigaDB [2 | This study |
| | *Isoetes tegetiformans* | ItBK1 | scaffold-PKOX-2003139 | kp [26], GigaDB [2 | This study |
| | *elaginella moellendor* | SmoBK1 | 441904 | 1.0 [38], Phytozon v12 [24] | mez-Porras, et al. [3 Sussmilch, et al. [31] |
| | | SmoBK2 | 406662 | | |
| | *elaginella stauntonia* | SsBK1a | scaffold-ZZOL-2011537 | kp [26], GigaDB [2 | This study |
| | | SsBK1b | scaffold-ZZOL-2003815 | | |
| | | SsBK2 | scaffold-ZZOL-2053187 | | |
| ants: ferns | *Salvinia cucullata* | ScBK1a | Sacu_v1.1_s0242.g02664 | 1.2 [39], FernBas | Sussmilch, et al. [31] |
| | | ScBK1b | Sacu_v1.1_s0055.g01445 | | |
| | | ScBK2 | Sacu_v1.1_s0171.g02440 | | |
| | *Azolla filiculoides* | AfBK1a | Azfi_s0018.g014960 | v1.2 [39], FernBas | Sussmilch, et al. [31] |

| Group | Species | Gene name | Accession | Sequence sourc | Gene reference |
|---|---|---|---|---|---|
| | | AfBK1b | Azfi_s0004.g008356 | | |
| | olypodium glycyrrhiz | PoglBK1 | scaffold-CJNT-2004894 | kp [26], GigaDB [2 | This study |
| | | PoglBK2 | scaffold-CJNT-2004895 | | |
| ants: gymnosperms | *Ginkgo biloba* | GbBK1 | Gb_05212 | GigaDB HiC [40] | This study |
| | *Gnetum montanum* | GmBK1 | TnS000904019t04 | DRYAD[41] | This study |
| | *seudotsuga menzies* (Douglas fir) | PmBK1 | PSME_06254 | .0 [42], PineRefS | This study |
| | *Picea abies* | PaBK1 | MA_10433576g0020 | 1.0, ConGenIE [4 | Sussmilch, et al. [31] |
| | *Picea glauca* | - | - | v1 [44], v3, v4 [45 Congenie | - |
| | *Pinus lambertiana* | - | - | .5 [46], PineRefS | - |
| | *Pinus pinaster* | PpiBK1 | PPI00062741 | Gymno PLAZA 1. [47] | This study |
| | *Pinus taeda* | - | - | .0 [48], PineRefS | - |
| ants: angiosperms | *Amborella trichopoda* | - | - | 1.0 [49], Phytozon v12 [24] | - |
| | *Arabidopsis thaliana* | - | - | TAIR10 [50], hytozome v12 [2 | - |
